# Ancient human genomes from Ladakh reveal Tibetan, South Asian, and Central Asian admixture over the last three millennia

**DOI:** 10.64898/2026.01.26.701789

**Authors:** Niraj Rai, Bhavna Ahlawat, Aparna Dwivedi, Snigdha Konar, Shristee Gupta, Esha Bandyopadhyay, Jose A. Urban Aragon, David Witonsky, Richa Rajpal, Prekshi Garg, Rashmi, Sachin Kumar, Chetan Vuppulury, Pankaj Singh Baghel, Shailesh Agrawal, Sonam Spalzin, Tashi Ldawa Thsangspa, Maanasa Raghavan

## Abstract

The trans-Himalayan region of Ladakh occupies a strategic position at the crossroads of South Asia, the Tibetan Plateau, and Central Asia, with archaeological evidence pointing to long-term cultural exchanges across these regions. However, the human genetic history of Ladakh remains largely unexplored. We generated paleogenomic data from seven individuals recovered from two sites in Western Ladakh - the Old Lady Spider Cave and Hanu - of which six are dated to 531-585 CE and one to the 19th century CE. The older individuals share substantial genetic ancestry with Tibetan groups but also harbor major contributions from two additional sources: one corresponding to the currently-oldest observation of the Ancestral North Indian genetic component that characterizes several present-day populations in North India and Pakistan, and another related to ancient Central Asian groups, with admixture events occurring between ∼2,100-2,500 years ago. In contrast, the later individual falls within a previously described ancient northern Himalayan genetic cline based on ∼1,100-1,300-year-old individuals from Himachal Pradesh, with ancestries related to ancient Tibetan and Steppe-related sources. Stable isotope analysis suggests that these individuals were local to Ladakh in late life and practiced an agro-pastoralist subsistence. Our study establishes that Ladakh’s central role in Eurasian economic and socio-cultural networks was shaped by dynamic and sustained gene flow linking high-altitude Himalayan groups with both lowland South Asia and Inner Asia.

## Introduction

The high-altitude cold desert of Ladakh, nestled in the trans-Himalayan region, is bound by the Karakoram Range in the north and the Zanskar Range in the south. Its rugged terrain consists of towering mountains, glacial systems, and deeply incised valleys, highlighting its impressive geological history. The hydrology and geomorphology of the region are primarily shaped by the Indus River and its tributaries, and its high-altitude passes have for long served as corridors for movements and exchanges, linking Ladakh to Central Asia, Tibet, and the rest of South Asia (1–5). These networks facilitated the movement of diverse commodities like pashmina wool, salt, textiles, tea, and spices, especially through the flourishing Silk Road, while concurrently acting as conduit for cultural and linguistic exchanges (6). The rich exchange of people and goods established Ladakh as an integral and dynamic zone of socio-economic interactions between South Asia, Central Asia, and the Tibetan Plateau.

Archaeological evidence from Ladakh, including lithic and bone tool assemblages, hearth features, and faunal remains as well as ancient monasteries, burial traditions, petroglyphs, and abandoned transcontinental trade routes, provide support for long-term human occupation and movements. While there is patchy evidence of inhabitation in the prehistoric period, material culture suggests early human groups in the Neolithic may have practiced transhumance and pastoralism, likely residing in seasonal camps (5, 7, 8). Furthermore, typological parallels in artifacts and burial practices with regions such as Swat (∼3000-500 BCE), Tibet (e.g., Phrang mgo, Gonggar), and Kinnaur in Himachal Pradesh (e.g., Kanam, ∼540 BCE) indicates prehistoric cultural interactions across the trans-Himalayan zone (9–11). Beyond the trans-Himalayan region, Ladakh’s cultural contacts with Central Asia and the Steppe region are evidenced in the ubiquitous rock art dating back to the Bronze and Iron Ages (second and first millennium BCE), depicting different stylistic elements such as animals and diverse ornamental and geometric line designs (12, 13).

Diffusion of cultural elements from Central Asia are additionally observed in the ceramic assemblages from the protohistoric and later periods (starting ∼3rd century BCE); Western Ladakh (Purig and Nubra) shares Central Asian technological and stylistic ceramic features such as the use of slow wheel and red/orange burnished slip finish, differentiating it from corded wares found in Eastern Ladakh (Changthang, Zanskar, Upper Ladakh) with links to Western Tibet and Spiti (14). Influences from neighboring northern regions of South Asia (15, 16) and Tibet (17), dating to early and late first millennium CE, respectively, can similarly be seen in rock inscriptions and art. In fact, North Indian, likely Kashmiri, architecture and art styles are observed in Buddhist monasteries and stupas dating to the 2nd century CE, hence pre-dating the Tibetan expansion, but also in later Tibetan-style monasteries from the 10th-11th centuries CE (18). Later architectural style, evident in the Leh Palace, was heavily influenced by the Tibetan architectural style, highlighting enduring political and cultural connection with Western Tibet (18). Chronicles, dating roughly between the 9th and 14th centuries, provide further accounts of contacts in ancient times (14, 19).

Unlike archaeological evidence, genetic studies on ancient and present-day humans in Ladakh are scarce. Few studies on uniparental and limited genome-wide markers report substantial and heterogeneous admixture in present-day Ladakhi populations from East, lowland South, and Central/West Asian sources (20–24), supporting dynamic movements into Ladakh over time. This genetic diversity is also reflected in the paleogenomic observations of ancestries related to Steppe and South and Central Asian groups in the neighboring regions of Spiti Valley and Kinnaur (Himachal Pradesh), dated to ∼1,300-1,100 years before present (yBP) (25), and Western Tibet, dated to ∼2,300-150 yBP (26, 27). Yet, the lack of ancient DNA data from Ladakh limits our understanding of this region’s biological connections with neighboring areas and of its past inhabitants.

This study presents genome-wide and isotope (carbon, nitrogen, and oxygen) data from ancient humans from Ladakh, dating between the mid-first millennium CE and the 19th century CE. The ancient individuals were sampled at two sites: Old Lady Spider Cave or Gachu Lhabrog (Kargil district) and Hanu (Leh district). Both sites are located in western Ladakh (Purig), with proximity to trade routes linking Ladakh to Central Asia and western China (Yarkand in present-day Xinjiang) via the Silk Road (28). There are no associated artifacts or contextual information from either site, making the genetic, radiocarbon dating, and isotope datasets generated in this study an invaluable window into Ladakh’s past. We report genetic ancestries related to ancient and present-day groups in Tibet, Central Asia/Steppe, and Northern regions of South Asia in these individuals, providing the first paleogenetic evidence in support of the dynamic socio-cultural and political interactions in ancient Ladakh as well as the currently-oldest observation of the ‘Ancestral North Indian’ (ANI) genetic component that characterizes several North Indian and Pakistani populations today.

## Results

### Genomic data summary

We generated shotgun sequencing data from 11 ancient individuals from two high-altitude sites (>∼2,700 meters above sea level (masl)) in Western Ladakh: Hanu (N = 3) and Old Lady Spider Cave (N = 8) (Figure 1A, Table S1). Of these screened individuals, two Hanu (HANU-2 and HANU-3) and five Old Lady Spider Cave (LD-01, LD-02, LD-11, LD-36, LD-TT) individuals were resequenced to higher coverage ranging between ∼0.11-0.85x, and, ultimately, these seven individuals were used in the presented population genetics analyses (Figure 1A, Table S1). The five individuals from Old Lady Spider Cave were radiocarbon dated to 531-585 calibrated (cal) CE, while one Hanu individual (HANU-2) was dated to 544 cal CE and the other (HANU-3) to 1833 cal CE. Characteristic cytosine deamination and fragmentation patterns were observed along with <3% mitochondrial DNA contamination levels, indicative of authentic ancient DNA in our dataset (Figure S1A-K, Table S1). Analyses using both BREADR (29) and KIN (30) identified a pair of first-degree relatives at Old Lady Spider Cave: LD-01 and LD-36 (Table S7A-B); being the lower covered individual of the related pair, LD-36 was subsequently excluded from further autosomal analyses except Principal Component Analysis (PCA). Finally, all libraries were trimmed to different lengths based on the extent of terminal damage to minimize potential biases in downstream analyses (see Methods).

**Figure 1.**
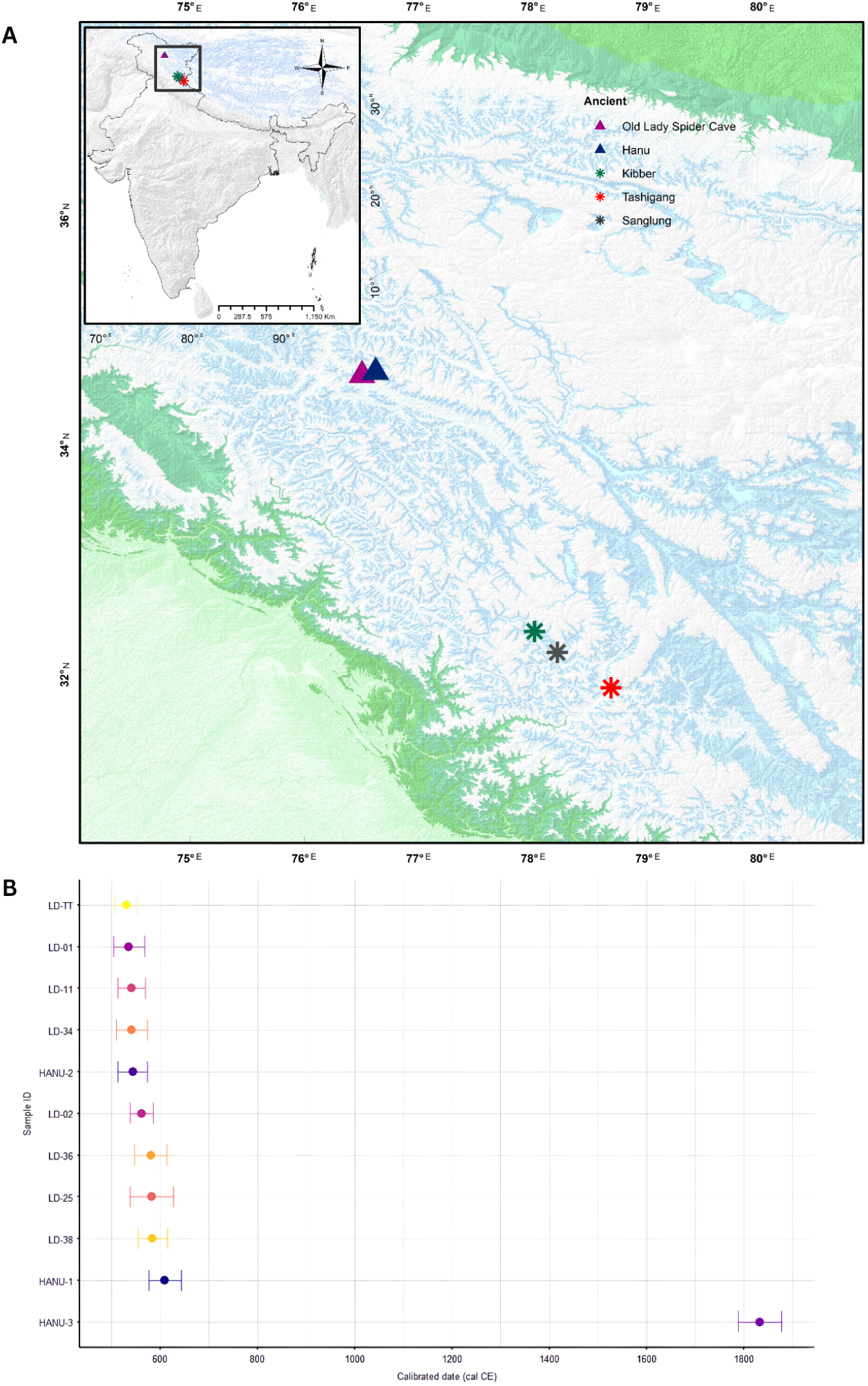
Spatio-temporal context of the sampled individual from Ladakh. A) Map of India with a zoom-in of Ladakh and Spiti (Himachal Pradesh) showing the locations of Hanu and Old Lady Spider Cave as well as sites from Himachal Pradesh with available paleogenomic data (from (25)), B) Radiocarbon dates (cal CE) for the ancient Ladakh individuals, calibrated with CALIB 8.2 using the IntCal20 Calibration curve. See Table S1 for more details.

### Genetic affinities and allele sharing patterns

On a PCA constructed with present-day Eurasian populations, the five Old Lady Spider Cave and two Hanu individuals fell along a genetic cline, intermediate between East and South Asians. Present-day members of this cline include populations that are primarily from the Northern Indian Himalayan region, including Ladakh and Himachal Pradesh: Brokpa, Changpa, Minero, Kashmiri Tibetans, and the HIM-A group from Lahaul and Spiti from (25) (Figure 2A, Figure S2A-B). At a finer scale, the individuals from Old Lady Spider Cave and HANU-2 clustered close to each other and a subset of present-day Brokpa and Changpa individuals from Ladakh (Figure 2A). In contrast, HANU-3, the youngest individual, clustered away from the Old Lady Spider Cave individuals, sharing higher genetic similarities with HIM-A, Kashmiri Tibetans, and ancient individuals from Lahaul and Spiti in Himachal Pradesh (Sanglung, Tashigang, and Kibber from(25) (Figure 2A). Overall, the ancient individuals from Ladakh show varying degrees of displacement on the PCA along the South and East Asian genetic cline, suggesting they likely have varying levels of genetic ancestries related to both sources.

**Figure 2.**
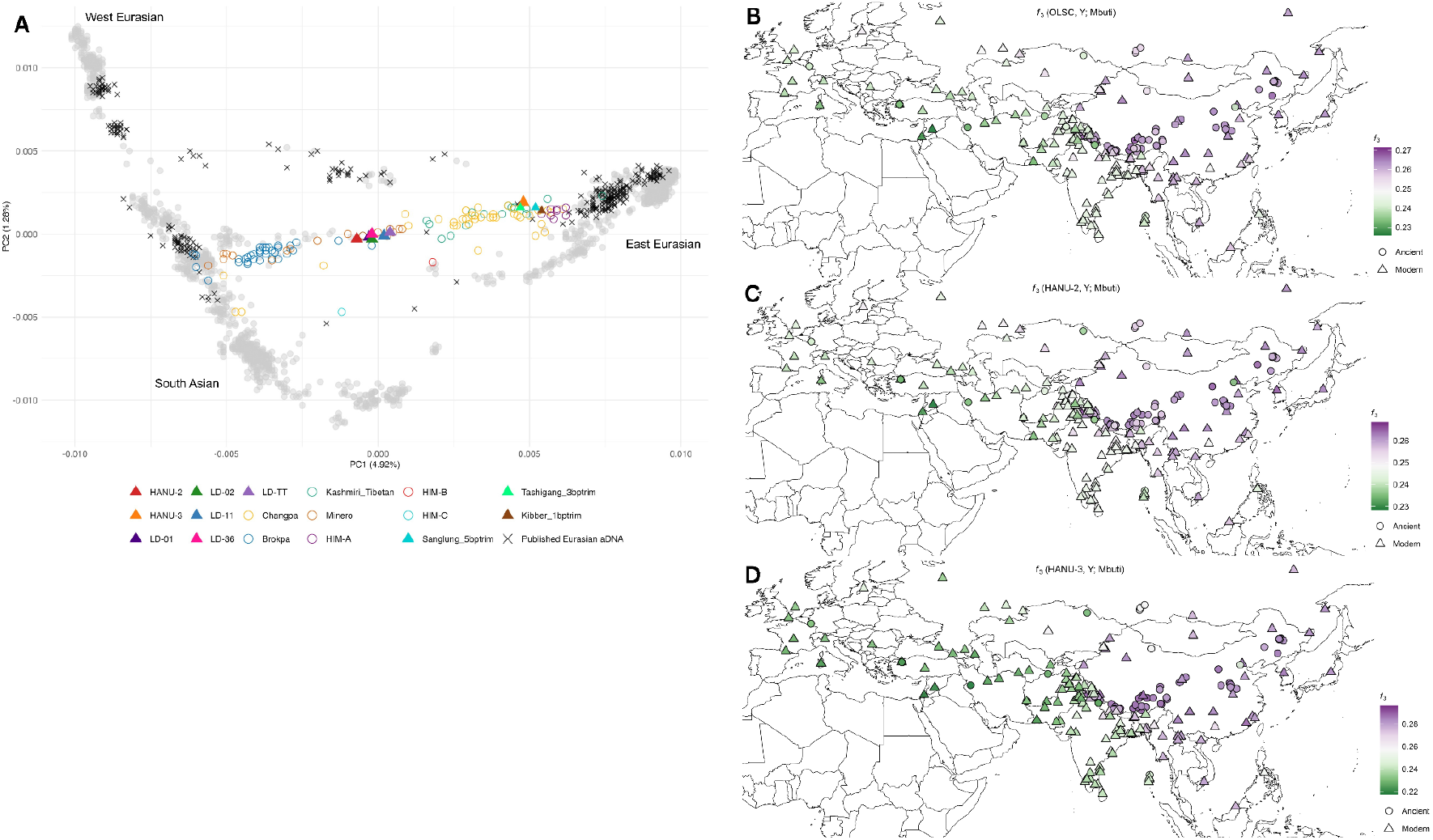
Broad-scale genetic affinities of the ancient Ladakh individuals. (A) Principal component analysis plot of the seven ancient Ladakh individuals projected onto a reference space built using present-day Eurasian populations. Published ancient and present-day individuals from Himachal Pradesh are also plotted. See Figures S2A-B for labeled plots. (B–D) Heat maps of outgroup-*f*_*3*_ statistics of the form *f*_*3*_(Test; Y, Mbuti), where Y represents diverse ancient and present-day Eurasian groups and Test represents the ancient Ladakh individuals: OLSC (B), HANU-2 (C), and HANU-3 (D). Color bars represent the *f*_*3*_ statistic value. See Tables 2A-C for more details.

Despite the PCA suggesting shared affinities to South and East Asia, outgroup *f*_*3*_-statistics revealed that all the ancient individuals from Ladakh shared higher genetic drift with ancient and modern groups from the Tibetan Plateau and Himalayas than with other South Asians (Figure 2B-D, Table S2A-C). Notably, HANU-3, which is shifted towards the East Asian cluster on the PCA, displayed higher relative drift sharing with East Asians than HANU-2 and OLSC (Old Lady Spider Cave group, comprising of LD-01, LD-02, LD-11, LD-TT). Higher allele sharing of ancient and present-day Western Eurasian groups with OLDSC and HANU-2 over HANU-3 and of Tibetan and East Asian groups with HANU-3 over HANU-2 and OLSC supports the latter sharing more genetic affinity with East Eurasians (Table S3).

Consistent with the above autosomal results, the ancient individuals sequenced in this study displayed a mix of maternal lineages reported in East, South, and Central Asia, as well as broader West Eurasia (Table S1). Due to the higher coverage of mitochondrial genomes, more individuals were included in this analysis compared to the autosomal analyses. Mitochondrial haplogroup (hg) C4a1, observed in the oldest individual sequenced in this study (LD-34), and in LD-25, has been reported in present-day individuals from Siberia, Mongolia, the Himalayas, and Central and West Asia (31–34). Hg U7a3b, observed in the next set of oldest individuals, LD-01 and LD-36, and hg U5a1a1 in LD-TT, have previously been identified in present-day populations from South Asia (i.e, Pakistan, Northern, and Central India), West Eurasia, Central Asia, the Near East, with hg U5a1a1 also reported in ancient individuals associated with the expansion of the Yamnaya culture (35–42). Haplogroups H2a1a and M52a1b, observed in the two Old Lady Spider Cave individuals (LD-02 and LD-11, respectively), have been reported in present-day individuals from South Asia (i.e., India, Nepal, and Pakistan), Central Asia (i.e., Mongolic-speaking populations), Southeast Asia, West Eurasia, North Africa, the Near East, and the Arabian Peninsula (43–46). Hg M65a, present in LD-38, is found in ancient and present-day individuals from Mesopotamia, Pakistan, Northwest India (i.e., Kashmir), Southern India, and Sri Lanka (35, 47–50). Finally, hgs T1a, F2g, and M9a1b, observed in HANU-1, HANU-2, and HANU-3, respectively, have been reported in present-day individuals from East and Central Asia (51, 52), West Eurasia, North Africa and the Arabian Peninsula (53–55), high-altitude populations from the Tibetan Plateau (52, 56–59), and, among Tibeto-Burman speakers in Northeast India (60).

### Admixture modeling and dating

Based on the differences in genetic affinities of the ancient Ladakh individuals to each other and to other Eurasian populations, we ran qpWave to assess whether these individuals are cladal with respect to a select set of reference groups (see Methods). We observed two cladal clusters: one consisting of the Old Lady Spider Cave individuals and HANU-2 and a second consisting of ancient and some present-day individuals from Himachal Pradesh (Sanglung, Tashigang, Kibber, and HIM-A) (Table S4). Notably, HANU-3 was only cladal with the Tashigang individual, which is consistent with their close placement on the PCA.

Based on the observed broad affinities of the ancient Ladakh individuals to populations from South Asia and the Tibetan Plateau, we began by attempting to model them as a mix of sources from these and neighboring regions. As one source, we used representative ancient and present-day groups from South Asia as well as ancient groups from Central Asia, West Asia, and the Steppe region. For the second source, we used ancient Tibetan groups from (26, 27, 61). We did not observe any working two source models for OLSC (Table S5A). In contrast, HANU-2 and HANU-3 yielded different two source models, in line with their variable genetic affinities to East and South Asians and, in turn, to each other. HANU-2 was modeled as a mix of ∼60% South Asian-related and ∼40% Tibetan-related sources (averages across all significant models) (Figure 3, Table S5B). The South Asian component in HANU-2 is best represented by present-day populations from North India and Pakistan with a higher proportion of the ANI-related genetic ancestry (62, 63). HANU-3 was modeled as ∼81% Tibetan-related and ∼19% Steppe-related sources (average of the top five significant models with Afanasievo as the Steppe-related source), with Afanasievo yielding consistently significant (p>0.05) and high p-values (Figure 3, Table S5C). The higher proportion of Tibetan-related ancestry in HANU-3 is concordant with the PCA and *f*-statistics results that illustrate this individual’s higher genetic affinity with Tibetan and other East Asian populations. Additionally, and consistent with the qpWave results, the ancestral sources and proportions in HANU-3 are similar to the individual from Tashigang and, more generally, to the other ancient Himachal and present-day HIM-A individuals modeled in (25).

**Figure 3.**
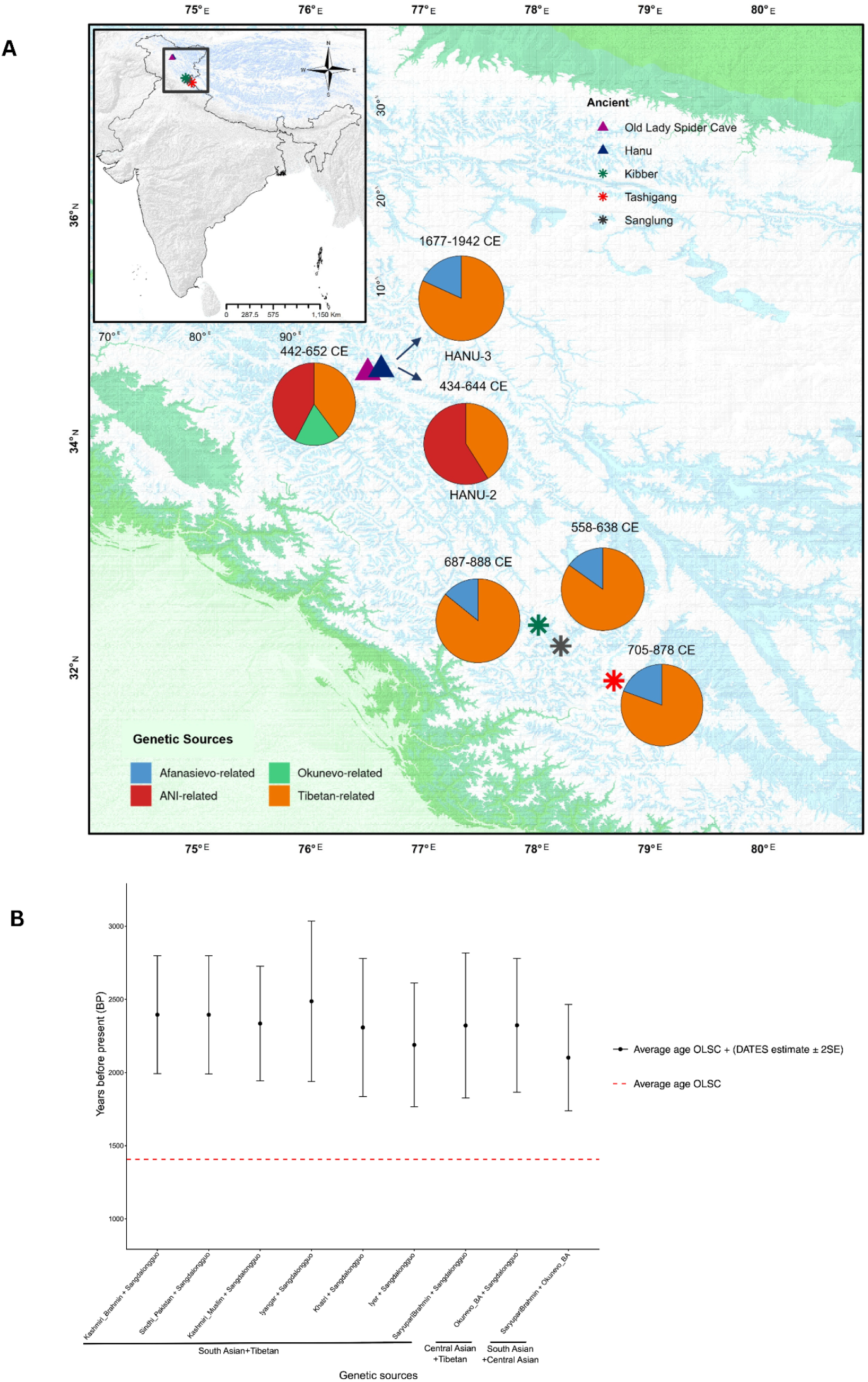
Admixture modeling and timings. (A) qpAdm-based genetic ancestry modeling of the ancient Ladakh individuals. Pie charts show inferred ancestry proportions for OLSC, the two Hanu individuals, and the ancient Himachal individuals Kibber, Tashigang, and Sanglung, modeled in (25). See Tables 5A-D for more details. Proportions of the genetic source proxies modeled in HANU-2, HANU-3, OLSC, Sanglung, Tashigang, and Kibber piecharts represent models with the highest p-values shown in Tables S5B-D and the supplementary material of (25). (B) Results of admixture time estimations from DATES for OLSC using combinations of genetic sources representing South Asian-, Tibetan-, and Central Asian-related ancestries, based on the top significant models in qpAdm (see Methods). Point estimates represent admixture dates in years before present ± 2SE for models passing the DATES significance criteria (see Methods), computed as 1407.5 years + (DATES estimate in generations × 30 years). Models are ordered from left to right by decreasing *Z* scores (Table S6). The red dashed line marks the average calibrated age of the OLSC group used in the analysis (1407.5 BP; Table S1).

Although OLSC failed to yield significant (p>0.05) models, we observed in the accompanying *f*_*4*_-statistics (‘gendstat’) that it consistently shared more drift with Okunevo_BA.SG (one of the reference set groups) than what could be explained by the two source models. Hence, we moved Okunevo_BA.SG as a third source and added Tarim_EMBA in the reference set to distinguish the Okunevo-related ancestry (64). We observed significant (p>0.05) three source models for OLSC, comprising ∼45% ANI-related, ∼17% Okunevo-related, and ∼38% Tibetan-related ancestries (averages across all significant models) (Figure 3, Table S5D). In light of this result, the inference of cladality between OLSC and HANU-2 in qpWave (Table S4) may result from reduced power due to smaller sample size since, despite sharing ANI- and Tibetan-related ancestries, a major differentiator between the two is the additional Okunevo-related ancestry in OLSC.

Timing of the admixture events reported above were inferred using DATES (65). We observed overall similar admixture dates for pairs of the three sources contributing to OLSC: ∼2,189-2,487 yBP for ANI- and Tibetan-related, ∼2,323 yBP for Okunevo- and Tibetan-related, and ∼2,102 yBP for Okunevo- and ANI-related (Table S6). We were unable to date the admixture event in HANU-2 and HANU-3 likely due to low coverage and sample size of one (Table S6).

### Paleodietary and late-life mobility reconstructions

Carbon (δ^13^C) and nitrogen (δ^15^N) stable isotope ratios were measured from bone collagen, which remodels throughout life and, thus, provides an integrated signal of individuals’ average adult diet (66–69). Most of the δ^15^N values in our data indicate moderate to high animal protein intake in the Old Lady Spider Cave and Hanu individuals, likely deriving from terrestrial sources such as sheep, cattle, or goat (Table S8, Figure S3A) (66, 69). The δ^13^C values for the majority of the individuals range between -18.0‰ to -19.1‰ (Table S8, Figure S3B), representing a C3 plant-based terrestrial ecosystem (68, 70, 71). These results support a pastoralist or agro-pastoralist subsistence, also prevalent among populations in the region today. While the tight clustering of isotopic values for most individuals within each site reveals a fairly homogeneous dietary regime, inter-site differences indicate potentially different mechanisms at play, including differences in the protein consumption or differential stress/disease status of individuals at the two sites.

Oxygen isotope (δ^18^O) composition was investigated to evaluate whether the individuals from the Old Lady Spider Cave, the site with more sampled individuals and modeled with diverse genetic ancestries, constituted local residents or migrants. The estimated isotope composition of ingested water-equivalent, or δ^18^O_dw_, based on bone bioapatite measurements (δ^18^O_c_) from the ancient individuals, ranged from -13.8‰ to -10‰ VSMOW (Table S9, Figure S5). Comparison with published δ^18^O values in precipitation from broader India revealed similar values in the Himalayan high-altitude region, including Ladakh, today (Figure 4A). While we cannot rule out early-life migrations not reflected in bone or migrations from a neighboring region with a similar hydrological range, our result does not exclude these individuals being local residents. While some (∼4‰ variance) intra-site variation in values is observed and may indicate individual-level mobility within the Ladakhi topography, minor seasonal changes, or measurement noise, the overall tight clustering of individuals and lack of clear outlying individuals, suggests this group primarily resided in a single high-elevation hydrological zone in the years represented by bone turnover.

**Figure 4.**
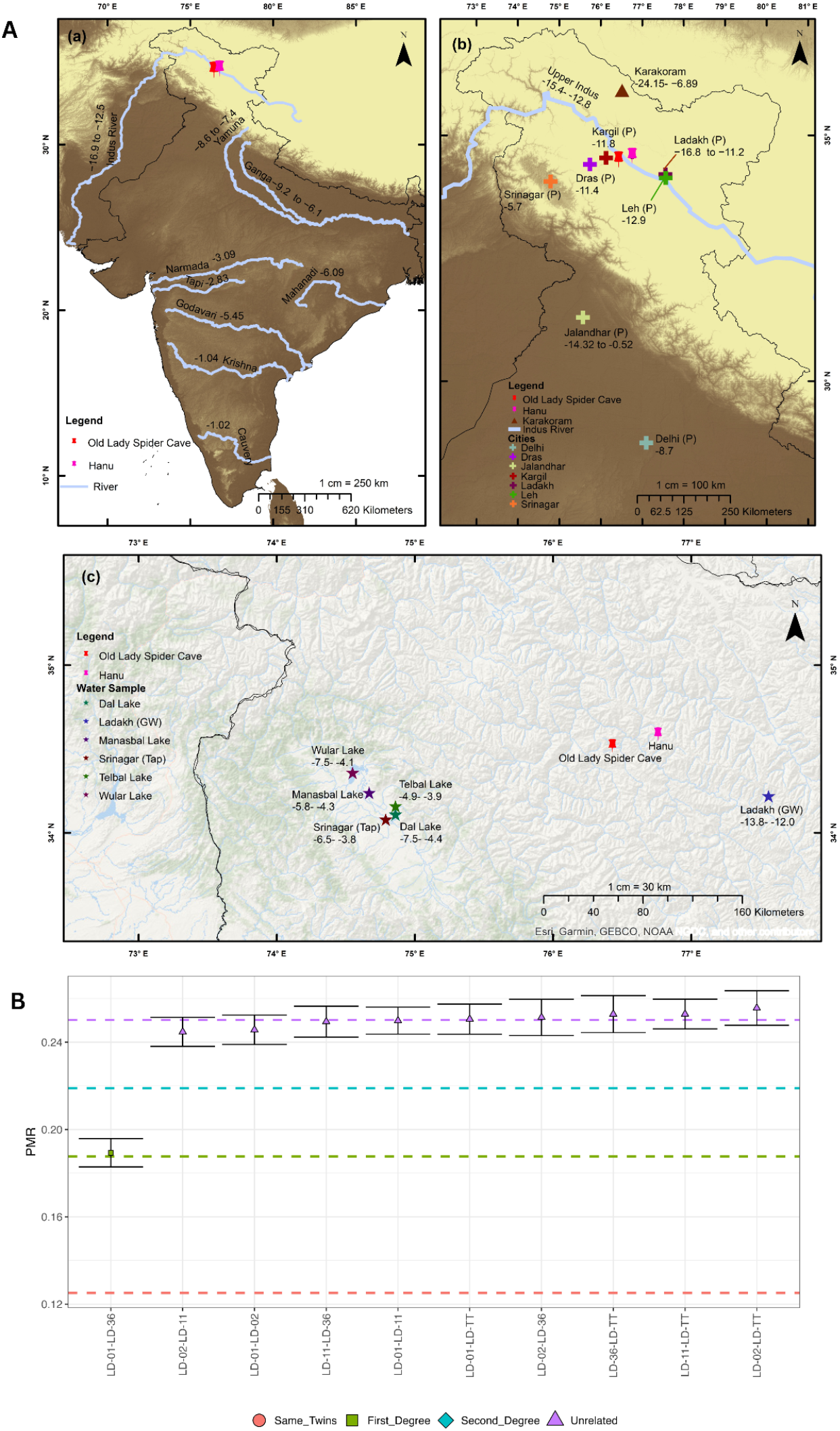
δ^18^O landscape of precipitation across India and biological relatedness between the ancient Ladakh individuals. A) Compilation of present-day oxygen isotope (δ^18^O) values from India and neighboring regions, providing a hydro-climatic baseline for archaeological interpretation. Values are reported in ‰ relative to VSMOW standard. (a) Spatial trend in river water δ^18^O values across India, illustrating a north-to-south isotopic gradient. (b) Point location data for precipitation (P) δ^18^O from different cities or locations in Northern India, supplemented by ice core data from the Karakoram range and Upper Indus River. (c) Zoom-in map of Kashmir and Ladakh regions illustrating δ^18^O values for groundwater (GW), tap water (Tap), and major lakes, compiled from (109–117). B) Pairwise mismatch rate (PMR)-based biological relatedness among the Old Lady Spider Cave individuals inferred using BREADR. Horizontal, colored dashed lines indicate expected PMR ranges for identical/twin, first-degree, second-degree, and unrelated relationships. See Table S7A for more details.

## Discussion

The geographical position of Ladakh at the crossroads of South, Central, and East Asia has shaped its pivotal role in human migrations, cultural exchanges, and genetic admixture. In addition to support from archaeology, genetic studies in present-day populations of Ladakh have reported diverse ancestries in the region today (20–24). Yet, we lack an understanding of how this critical spatial positioning of Ladakh impacted its past genetic landscape, the temporal context to human movements into the region, and South Asia’s broader inter-connectedness with Inner Asia. Our study fills this critical gap by presenting seven ancient human genomes from two sites in Western Ladakh, spanning ∼530-1,800 CE, and complementing this with stable isotope data to understand past mobility and paleodietary patterns.

Our paleogenetic results highlight high levels of genetic diversity in the studied time period. Although all analyzed individuals show high genetic similarity with ancient and present-day Tibetan Plateau groups (Figure 2B-D), several notable differences in the non-Tibetan genetic ancestries emerge. Our analyses show the presence of two distinctly admixed groups dating to the mid-first millennium CE, one at the Old Lady Spider Cave (four individuals) and another at Hanu (one individual). At both sites, we observed substantial proportions of a genetic component related to ANI: ∼60% in the Hanu individual and ∼45% in the Old Lady Spider Cave individuals (Figure 3). ANI is a statistically-constructed group that was likely formed sometime after 4,000 yBP and contributed genetic ancestry to several present-day Indo-European speakers in South Asia, primarily living in northern India and Pakistan (62, 63, 72). In fact, the best-fitting ANI-related sources in our qpAdm models are present-day populations from these regions. Our observation of this genetic ancestry in the ancient individuals from Ladakh has more profound implications for the population history of South Asia as this constitutes the first and thus-far oldest empirical evidence of ANI-related genetic ancestry in the region. Moreover, the admixture timing of this component is ∼2,100-2,500 yBP in the Old Lady Spider Cave individuals (Table S6). These results imply that ANI-related groups were present in the region by at least ∼2,100-2,500 yBP and contributed to the genetic ancestries of diverse groups in high-altitude Ladakh, in addition to low-altitude ancestors of many present-day northern South Asians. Notably, this admixture event seems to differ from lowland South Asian admixture in a present-day northern Himalayan individual (HIM-B) from Himachal Pradesh, dated to ∼2,400-3,000 yBP, in that the source of the ancestry in HIM-B is best approximated by present-day populations that place intermediately on the ANI-Ancestral South Indian (ASI) cline (25). This suggests a heterogenous landscape of ancient admixture events between high-altitude Himalayan and diverse lowland South Asian groups along the ANI-ASI cline.

Our inferences are supported by the archaeological record. By the early first millennium CE, the appearance of rock inscriptions and scripts in ancient Indic scripts, such as *Kharo*ṣṭ*h*ī and *Br*ā*hm*ī, suggest cultural influences from neighboring northern regions of South Asia (15, 16). Likewise, the oldest Buddhist stupa in Ladakh, the Sani Kanika Stupa in Zanskar, is proposed to have been originally constructed in the 2nd century CE during the Kushan period and thought to reflect early North Indian architectural style (18, 73). As such, our results imply that at least some of these interactions transcended cultural exchanges and resulted in genetic admixture, though our data is unable to identify whether this genetic mixing may have happened in Ladakh or elsewhere.

Although these ancient study individuals share genetic ancestries related to both ancient Tibetans and ANI, the Old Lady Spider Cave individuals additionally have genetic ancestry related to individuals affiliated with the Early-Middle Bronze Age Okunevo culture from the Altai region (Figure 3), dated to ∼2,000-2,500 yBP. Several rock art motifs in Ladakh, including mascoids and hunting scenes depicting stylized deer and ibex, are linked to the Bronze Age Okunevo and Andronovo cultures and later Iron Age cultures such as Saka, South Siberia, Altai, Mongolia, and Tuva (12, 13). While the cultural ties with Central Asia and the Steppe region are proposed to date back to the Bronze Age, the admixture date for this ancestry in the ancient study individuals is ∼2,100-2,300 yBP (Table S6), which represents the Iron Age rather than the Bronze Age. This indicates that the true source may not be Okunevo, but rather a younger group to which Okunevo or related groups contributed genetic ancestry. This scenario would also be more parsimonious given the distance between Ladakh and the Okunevo geographical range in South Siberia, and Central Asia being a more proximal corridor into Ladakh and surrounding regions. Expanded paleogenetic studies from Ladakh and Central Asia should investigate the true source of this ancestry in the Old Lady Spider Cave individuals.

Our estimates of the admixture dates for both the ANI- and Okunevo-related ancestries overlap, suggesting rapid genetic exchanges between these and the Tibetan sources (Table S6), though the exact spatial context to these events needs to be addressed in future investigations. Interestingly, Tibetan cultural influence in Ladakhi rock art dates back to the second half of the first millennium CE, with westward expansions starting with the Tibetan Empire between the seventh and eleventh centuries CE (17) and again around 10th-11th centuries CE, linked to the expansion of Tibetan Buddhism (18). Clearly, our admixture dates pre-date these known expansions into Ladakh, suggesting that at least some of these admixture events may have happened in neighboring areas or that more ancient evidence of Tibetan contacts are yet to be uncovered in Ladakh.

The youngest individual in our dataset from Hanu (HANU-3), dated to ∼1,800 CE, was modeled as a mixture of predominantly Tibetan- and minor Afanasievo-related ancestries, similar to ancient (Sanglung, Tashigang, and Kibber) and some present-day (HIM-A) individuals from Himachal (Figure 3). Close genetic similarity in several analyses between this ancient Ladakh and Himachal individuals illustrates that the ancient Himachal lineage (25) was present more widely in the northern Himalayas, at least in historical times.

Our genetic and isotope data also enabled us to make individual-level inferences, including paleodiet, mobility, and genetic relatedness. The oxygen isotope analysis on an expanded set of Old Lady Spider Cave individuals suggests that they were likely local residents. We, however, caution that technical (use of bone that remodels during an individual’s lifetime) and analytical (use of a regression equation derived from tooth carbonate on bone bioapatite and equivalence of precipitation to ingested water) limits a more conclusive inference. The paleodietary reconstructions, highlighting moderate to high animal protein and C3 plant consumption, supports a pastoralist subsistence that is in line with the local geophysical and ecological environment as well as present-day practices in the villages close to the sampled sites. Finally, although the lack of associated artifacts and fine-scale archaeological context hinders our understanding of the socio-cultural aspects of the studied individuals’ lives, we were able to reconstruct a partial family unit using biological relatedness as a measure. We identified a pair of first-degree genetic relatives at the Old Lady Spider Cave using BREADR (29) and KIN (30): LD-01 and LD-36 (Figure 4B, Table S7). Given that both share the same mtDNA hg (U7a3b), LD-01 is genetically male and LD-36 is genetically female, and LD-01 lived earlier than LD-36 based on their radiocarbon dates (Table S1), the most parsimonious inferred genetic relationship is siblings, also estimated by KIN (Table S7B). It is intriguing that we found a pair of close relatives among the five analyzed individuals at this site. Along with the oxygen isotope results, the inference of related individuals at the site adds further support for them likely being local residents rather than traders who may have been passing through the region.

This study contributes to our limited but growing understanding of human migratory histories in the northern Himalayas. Considering the genetic diversity and admixture times we observe, the Silk Road, but possibly also older trade networks, are likely to have played a crucial role in facilitating movement of people across Tibet, South Asia, and Central Asia, along with goods like gold, borax, salt, and pashmina (6). Our work illuminates a central role for Ladakh and South Asia within the broader Eurasian socio-cultural and economic sphere and provides a spatio-temporal snapshot of the ANI-related ancestry in the past.

## Methods

### Site descriptions, sampling, ethics, and community engagement

The studied sites in Western Ladakh lack associated cultural artifacts and, hence, thorough contextual characterization. As noted below, sampling at these sites derived from an exploratory survey and accidental finding. Sampling and scientific analyses of the human bones were conducted with appropriate approvals (see descriptions below) and oversight from the institutional review board of Birbal Sahni Institute of Palaeosciences (BSIP), Department of Science and Technology, Ministry of Science and Technology, Government of India (BSIP/Ethical Approval/2021/Letter-1).

Old Lady Spider Cave (34.5517°N, 76.4241°E), also Gachu Lhabrog, is a high-altitude grazing patch, situated at an altitude of 4,000 masl, ∼3.5 kilometers from the village of Yokma Kharbu in Ladakh. The cave is complex in structure with a total of three chambers linked by narrow galleries. The three chambers of the cave include cist burials, petroglyphs, and partially mummified human and faunal remains. The innermost chamber is only accessible by crawling and has remained largely undisturbed. Along with skeletal remains, the cave also has ceramic fragments and premodern beads made of stone, corals, and glass. The cave has remained openly accessible for long and was likely a target of looting over time. This may also explain the scattered nature of the human bones across the cave. While we lack cultural context at this site, the tight distribution of radiocarbon dates (Figure 1B, Table S1) indicates that the site may have plausibly been in use for a short period, after which its use was abandoned by local residents who may have seeked out alternative funerary practices. The ancient human bones were collected as part of a surface exploration effort in 2019 led by author S.S. (Archaeological Survey of India) along with N.R. who received approval from BSIP.

The Hanu site/mound is located in the village of Yokma Hanu (34.5906°N, 76.6218°E) in the Khaltsi tehsil of Leh district, at an altitude of 2,760 masl. It lies between Chorbat Pass and the Indus Valley. Historically, Hanu lay on an ancient route connecting Baltistan with Ladakh, with the Chorbat Pass considered as a natural boundary between these two regions. The village is fortified, indicating that they may have experienced raids from Baltistan in the past. Today, it is predominantly settled by members of the Dardic-speaking Brokpa tribe. The ancient individuals analyzed in this study were originally given proper burials, but accidentally unearthed in 2022 during the ‘Jal Jeevan Mission’, a water provision scheme by the Government of India. They have been curated since then by the local district administration that granted permission for the analyses presented in this study. There were no associated artifacts found in the Hanu burials.

Community engagement sessions to discuss the results are planned in villages close to both sites and will be conducted concurrently to the review process.

### Ancient DNA data generation

All data generation work was carried out in the ancient DNA laboratory in Birbal Sahni Institute of Palaeosciences, Lucknow, India. Prior to processing, the samples were surface-wiped with 2% bleach, followed by millQ water, and further UV-treated to minimize surface contamination. Next, approximately 3mm of outer layer was removed using a drill machine, and 30-50 mg of fine powder was generated for DNA extractions using the lowest drill setting. DNA extractions were carried out using the silica bead method with an improved binding buffer described in (74). Double-stranded libraries were prepared from each extract for an initial round of screening, following the protocol in (75). The libraries were sequenced on Illumina NovaSeq 6000 in 100 bp single-read mode. Seven individuals (LD-01, LD-02, LD-11, LD-36, LD-TT, HANU-2, and HANU-3) were subsequently sequenced to higher coverage and used in the analyses described below.

### Data processing, contamination estimation, and genotype calling

AdapterRemoval v2.5.3 (76) was used to remove residual sequencing adapters, consecutive Ns, low-quality bases (`--minquality=20`), and reads shorter than 30 bp. Reads were then mapped to the human reference genome GRCh37 and the mitochondrial revised Cambridge Reference Sequence (rCRS) build 17 using Burrows-Wheeler aligner (BWA) v0.7.17 (77). Reads were subsequently filtered for a mapping quality score of >30 using SAMtools v1.18 (78). Duplicates were removed with Picard MarkDuplicates v2.26.10 (79), and local realignment was carried out using GATK v3.8 (80). Mismatches and deleted reference bases (MD) tags were added using SAMtools calmd(78). Ancient DNA damage patterns were obtained with mapDamage v2.0 (81). Molecular sex was determined using the method described in (82). mtDNA contamination estimates were obtained using the R package contamMix (R v4.2.1, contamMix v1.0-10) (83). The autosomal, X, Y, and mtDNA coverages were estimated using SAMtools coverage (84).

To reduce the effect of post-mortem damage while generating pseudohaploid calls, end-trimmed bam versions of the seven ancient individuals were generated based on their mapDamage profiles (cytosine deamination). The trimBam module of BamUtil v1.0.14(67) was used to trim 3 bp at the 5’ and 3’ ends of the reads for the five Old Lady Spider Cave individuals (LD-01, LD-02, LD-11, LD-36, and LD-TT), 4 bp for HANU-2, and 7 bp for HANU-3.

For genotype calling, a pileup file was created using SAMtools mpileup v1.17 (65) with “`-R`“ and “`-B`“ flags. Pseudohaploid genotypes were generated using a minimum base and mapping quality threshold of 20 and 30, respectively, with parameter `-randomHaploid` using the pileupCaller program in the package sequenceTools v4.0.5 (66).

### Compilation of the dataset

The newly generated data from this study was merged with the Allen Ancient DNA Resource (AADR v54.1.p1) genotyped at ∼1.23 million sites (85), whole-genome sequencing data from present-day individuals in the GenomeAsia 100K dataset (86), ancient DNA data from the Tibetan Plateau genotyped at ∼1.23 million sites (26, 27), ancient and present-day whole-genome sequencing data from the northern Himalayas (25), whole-genome sequencing data from present-day individuals from the Tibetan Plateau (87–89), and genotyping data from present-day individuals from Kashmir and Ladakh at ∼643k sites on the Illumina HumanOmniExpress-24-v1 array (23). Individuals flagged as ignored, low-coverage, contaminated, or as a relative in AADR were filtered out. The resulting dataset was further filtered to remove individuals with greater than 90% missing data. The final dataset retained a total of 1,109,329 sites.

### Mitochondrial haplogroup analyses

mtDNA reads were extracted from the bam files of the ancient individuals from this study. Variant calling was done using bcftools mpileup v1.19 and bcftools call (90) haplogroups with Haplogrep 3 v3.2.1 (91).

### Principal Component Analysis

Principal component analysis (PCA) was performed using the program smartpca v16000 from the package EIGENSOFT v7.2.1 (92, 93) with parameters, `lsqproject: YES`, and `shrinkmode: YES`. Present-day groups from South Asia (including those from Ladakh, Himachal Pradesh, and Kashmir), the Tibetan Plateau, Central Asia, West Asia, Europe, East Asia, and Southeast Asia were used to construct the principal components (PC) space.

### *f*-statistics

Outgroup-*f*_3_ statistics were computed using the *qp3pop* program in the R package, ADMIXTOOLS 2 (94), with default parameters. The tested topology, *f*_3_(Mbuti; Test, Y), quantified shared genetic drift between a Test group and Y populations representing diverse ancient and present-day Eurasian groups. Test groups are OLSC (Old Lady Spider Cave group, comprising LD-01, LD-02, LD-11, LD-TT), HANU-2, and HANU-3.

*D-*statistics were computed using the *qpdstat* program in the R package, ADMIXTOOLS 2 (94). We evaluated topologies of the form *D*(OLSC/HANU-2/HANU-3, HANU-2/HANU-3; Eurasians, Mbuti), to test allele sharing patterns of a subset of diverse East and West Eurasians with pairs of the sequenced ancient individuals from Ladakh.

All outgroup-*f*_3_ and *D*-statistic tests involving fewer than 30,000 SNPs were excluded.

### Estimating cladality and admixture proportions

The qpWave program in the R package ADMIXTOOLS 2 (94) was used to test whether pairs of the ancient Ladakh individuals, as well as the OLSC group, are cladal relative to a set of reference populations used in (25): Mbuti.DG, China_AR_EN, Fujian_EN, Shandong_EN, Guangxi_EN, Okunevo_BA.SG, Zongri5.1k, Chamdo2.7k, Shannan3k, GBSL_old, Afanasievo, Onge, Russia_MLBA_Sintashta.SG. The qpAdm program from the same package was to estimate ancestry mixture proportions for OLSC, HANU-2, and HANU-3. For two-source models, the reference population listed above for qpWave was used. For three-source models for OLSC as the target, Tarim_EMBA1 was added to the reference set in place of Okunevo_BA.SG that was used as one of the three sources. Analyses were run with the parameter allsnps=TRUE.

### Dating genetic admixture events

To estimate the timing of admixture events among the ANI-related, Tibetan-related, and Steppe-related (Okunevo) genetic sources in the ancient Ladakh individuals, DATES v4010 (65) was used with parameters binsize: 0.001, maxdis: 0.9, jackknife: YES, qbin: 10, runfit: YES, afffit: YES, and lovafit: 0.45. Models were considered significant if the *Z*-score was ≥ 2, the mean (λ) < 200 generations, and the normalized mean square deviation (nrmsd) < 0.7 (65). Admixture time in yBP was estimated assuming 30 years as one generation. Source pairs were selected based on the best-fit qpAdm admixture models for each target, with a few representatives each of the broader ANI-, Tibetan-, and Steppe-related ancestries, in addition to Okunevo as a third source for OLSC (Tables 5B-D).

### Relatedness analysis

Genetic relatedness between pairs of the Old Lady Spider Cave was assessed using the R package BREADR (v1.0.0) (29) and KIN (30).

### Stable isotope analysis for Carbon (δ^13^C) and Nitrogen (δ^15^N) and radiocarbon dating

Sample preparation for stable isotope analysis and radiocarbon dating followed standard protocols (95, 96). Briefly, bone fragments from Old Lady Spider Cave and Hanu were demineralized in 0.5M hydrochloric acid at 4°C, rinsed with sodium hydroxide and water, and gelatinized. The resulting solution was filtered and purified collagen was lyophilized. Stable carbon (δ^13^C) and nitrogen (δ^15^N) isotope ratios were measured at the stable isotopes facility in Birbal Sahni Institute of Palaeosciences (BSIP) using Elemental Analyzer coupled to an Isotope Ratio Mass Spectrometer (EA-IRMS).

For radiocarbon dating, all collagen samples passing quality control were weighed and sealed into tin boats before introducing them into the elemental analyzer (EA) to undergo flash combustion. The evolved gases were purified and CO_2_ and N_2_ were separated using a gas chromatographic column and quantified using a thermally coupled detector. The purified CO_2_ was then transferred to the graphitization unit to prepare graphite targets for accelerator mass spectrometry (AMS) radiocarbon analysis. Reduction of CO_2_ to graphite was carried with H_2_ at 550°C within individual reactors to produce pellets of graphite. Graphitization was carried out using the Automated Graphitization Equipment 3 (Ionplus AG, Switzerland) coupled to a Vario Isotope Select elemental analyzer (Elementar Analysensysteme GmbH, Germany) and the graphite targets were measured at the Inter-University Accelerator Centre (IUAC), New Delhi, India.

Calibration was performed with CALIB 8.2 using the IntCal20 Calibration curve (97, 98). All the samples fell within the accepted archaeological C/N range ∼2.9 to ∼3.6, confirming good collagen preservation and its reliability for isotopic palaeodietary reconstruction and radiocarbon dating (Table S8, Figure S4) (99– 103).

### Stable isotope analysis for Carbon (δ^13^C) and Oxygen (δ^18^O)

Bone fragments of different individuals from Old Lady Spider Cave were cleaned mechanically and ultrasonically using Milli-Q water. Powdered samples were obtained using a diamond-tipped drill. To isolate bioapatite and remove diagenetic carbonates and organic contaminants, a standardized chemical pretreatment protocol was followed (104–106). Stable carbon (δ^13^C) and oxygen (δ^18^O) isotope ratios were measured on the purified bioapatite using isotope ratio mass spectrometer (IRMS) at the Birbal Sahni Institute of Palaeosciences, Lucknow, India (106). Isotope results are reported in standard delta (δ) notation in per mil (‰). Carbon isotope ratios are reported relative to Vienna Pee Dee Belemnite (VPDB). Oxygen isotope ratios are reported relative to VPDB and were converted to Vienna Standard Mean Ocean Water (VSMOW) scale using the equation: δ^18^O_VSMOW_= 1.3086 * δ^18^O_VPDB_ + 30.92 (107).

### Calculations and baseline data for Oxygen (δ^18^O) analysis

Ladakh lacks oxygen isotopic (δ^18^O) data from faunal samples of the same time period, which are often used to create a local isotopic baseline since animals often drink the same water as humans. An alternate and less direct approach to understanding mobility and paleoclimate of the region is through geological/ hydrological data since there is a high chance that humans directly consumed river water or stored precipitated water in the past (106). This can be used to construct spatial isoscape maps and paleoclimate models. The Old Lady Spider cave is situated in the western Himalayas, a region characterized by a cold, arid, and high-altitude climate. This environment produces meteoric water that is strongly depleted in the heavy isotopes ^18^O. The regional precipitation δ^18^O value of western Himalaya is -10.2‰ (108). Ladakh’s precipitation (P) δ^18^O value is -16.8‰ to -11.2‰ VSMOW (109, 110). The values of Upper Indus River water and Ladakh groundwater (GW) are -15.4‰ to -12.8‰ and -13.8‰ to -12.0‰ VSMOW, respectively (Figure 4A) (111–117). To interpret human mobility, the oxygen isotope composition of ingested water-equivalent (δ^18^O_dw_) was estimated from bone bioapatite (δ^18^O_c_) using an empirically derived carbonate-water regression: δ^18^O_c_ = 0.77 * δ^18^O_dw_ + 28.1‰ (118). Statistical analysis and data visualization were performed using R software (v2025.09.0+387).

## Supporting information

Large Supplementary Tables

## Data, Materials, and Software Availability

Data associated with this project is available on the European Nucleotide Archive Project, under accession number XYZ.

## Acknowledgments and funding sources

The authors are grateful to the Archaeological Survey of India, Mini Circle, Leh, for supporting this project. They are also thankful to Yountan Gyatso (Thorhasli Students Federation, Hanu) for facilitating meetings and conversations in Hanu for return of results. They acknowledge Birbal Sahni Institute of Palaeosciences institutional funds, project number 7.2 (to N.R.) and National Institutes of Health grant R35GM143094 (to M.R.).

