## Supplementary material for "Ancient human genomes from Ladakh reveal Tibetan, South Asian, and Central Asian admixture over the last three millennia": Supplemetary Figures and Tables

### Supplementary Tables and Figures

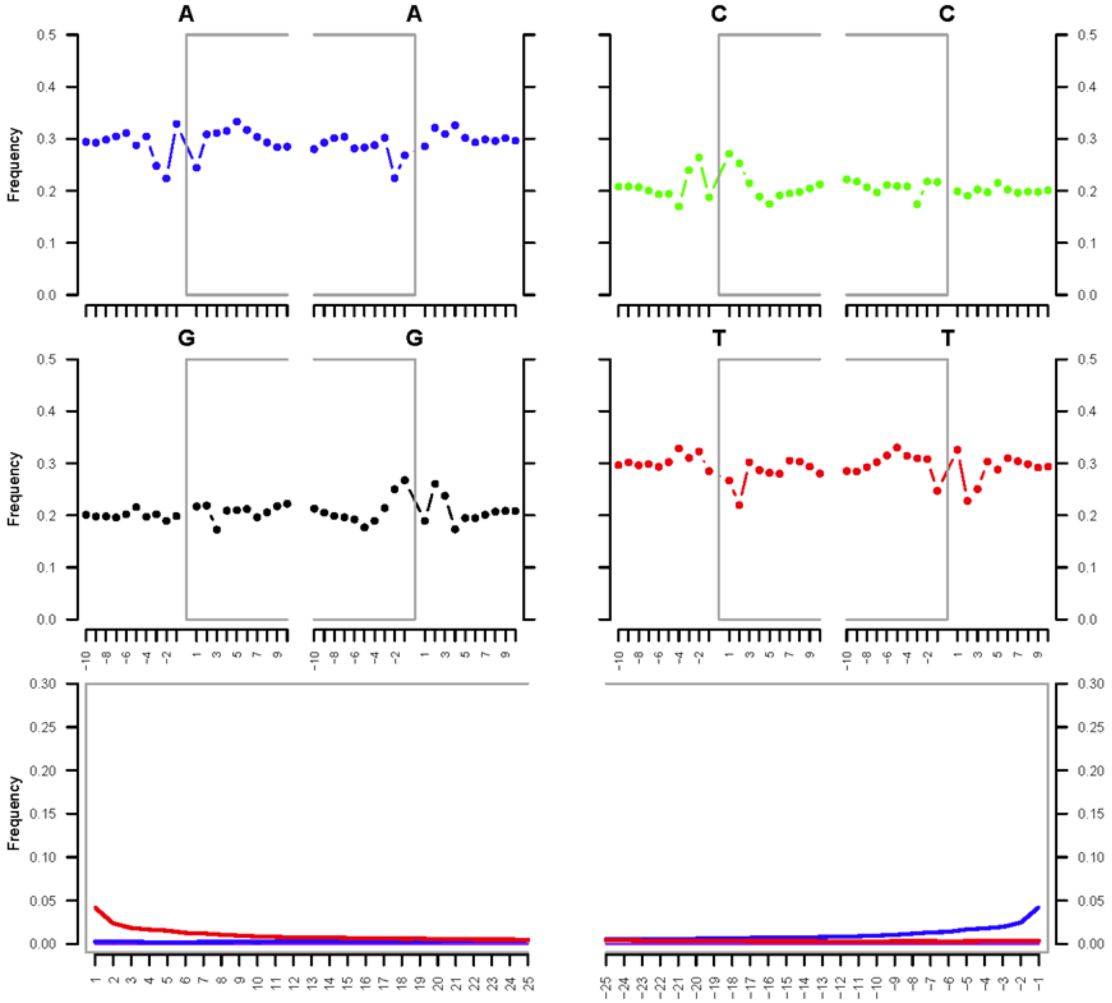


A. LD_1

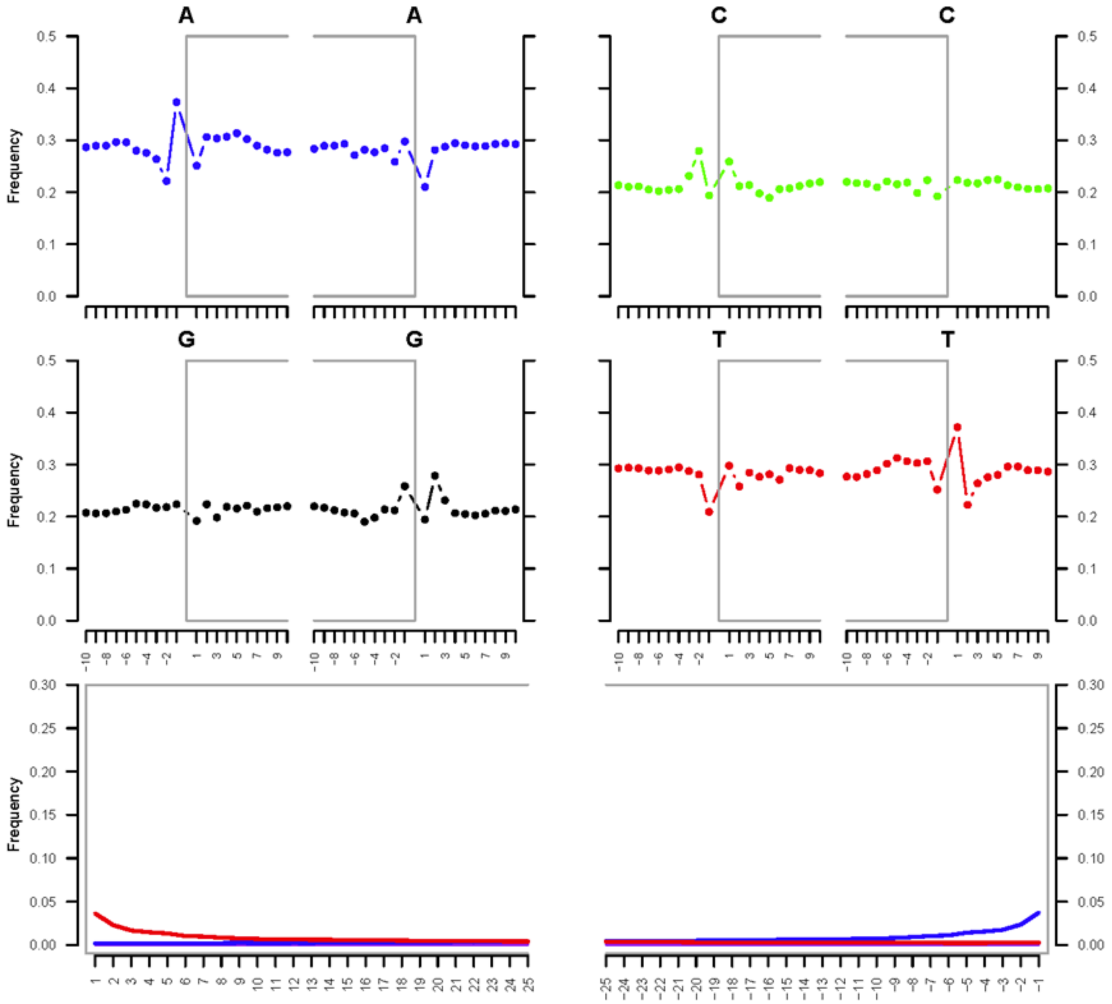


B. LD_2

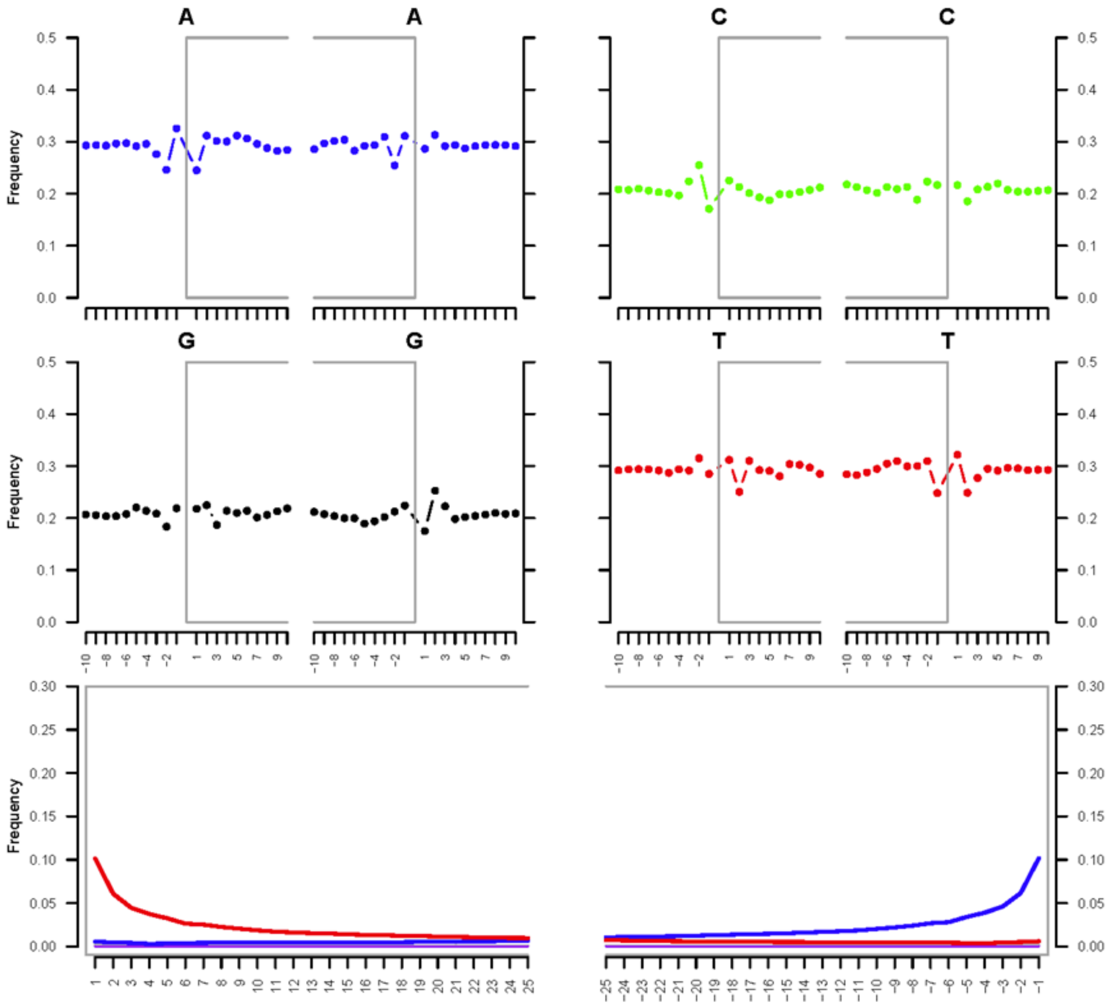


C. LD_11

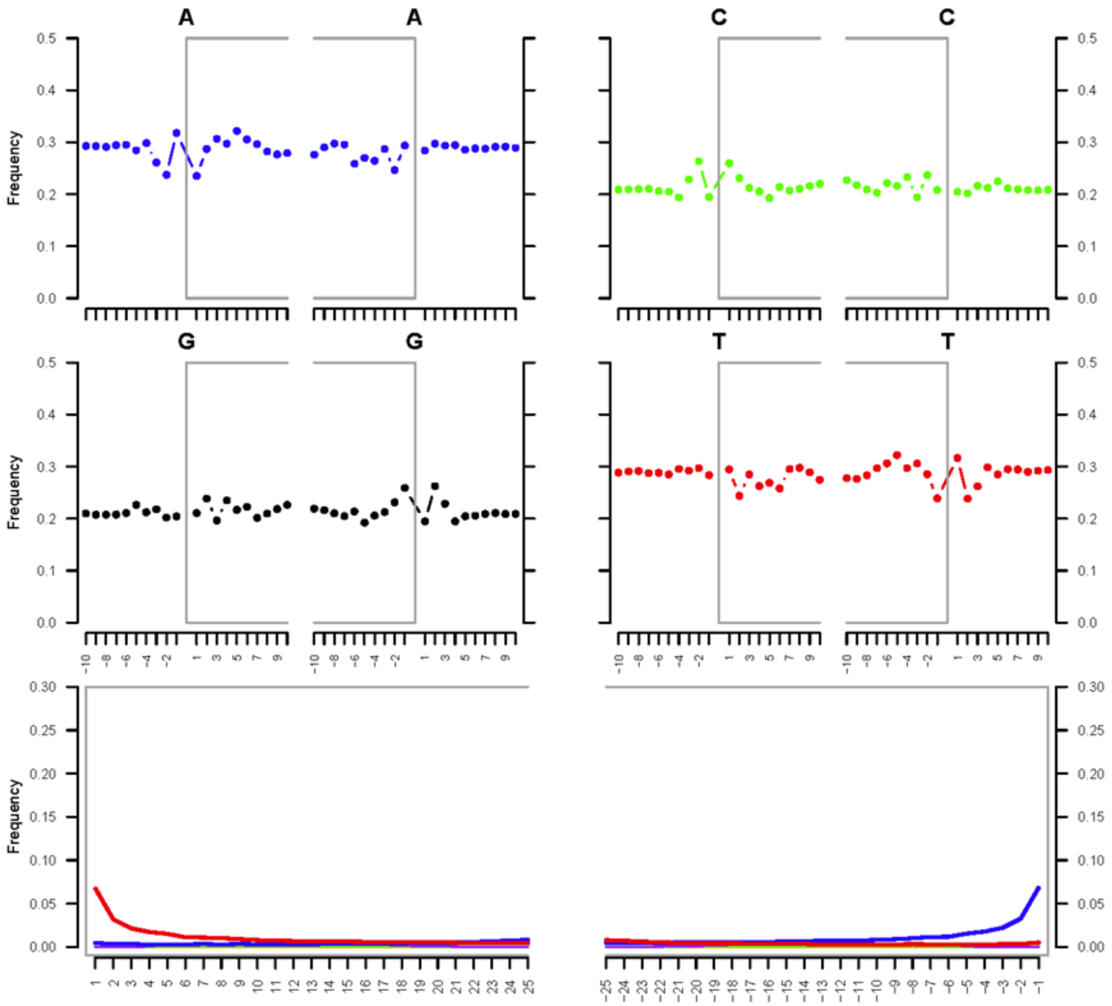


D. LD_25

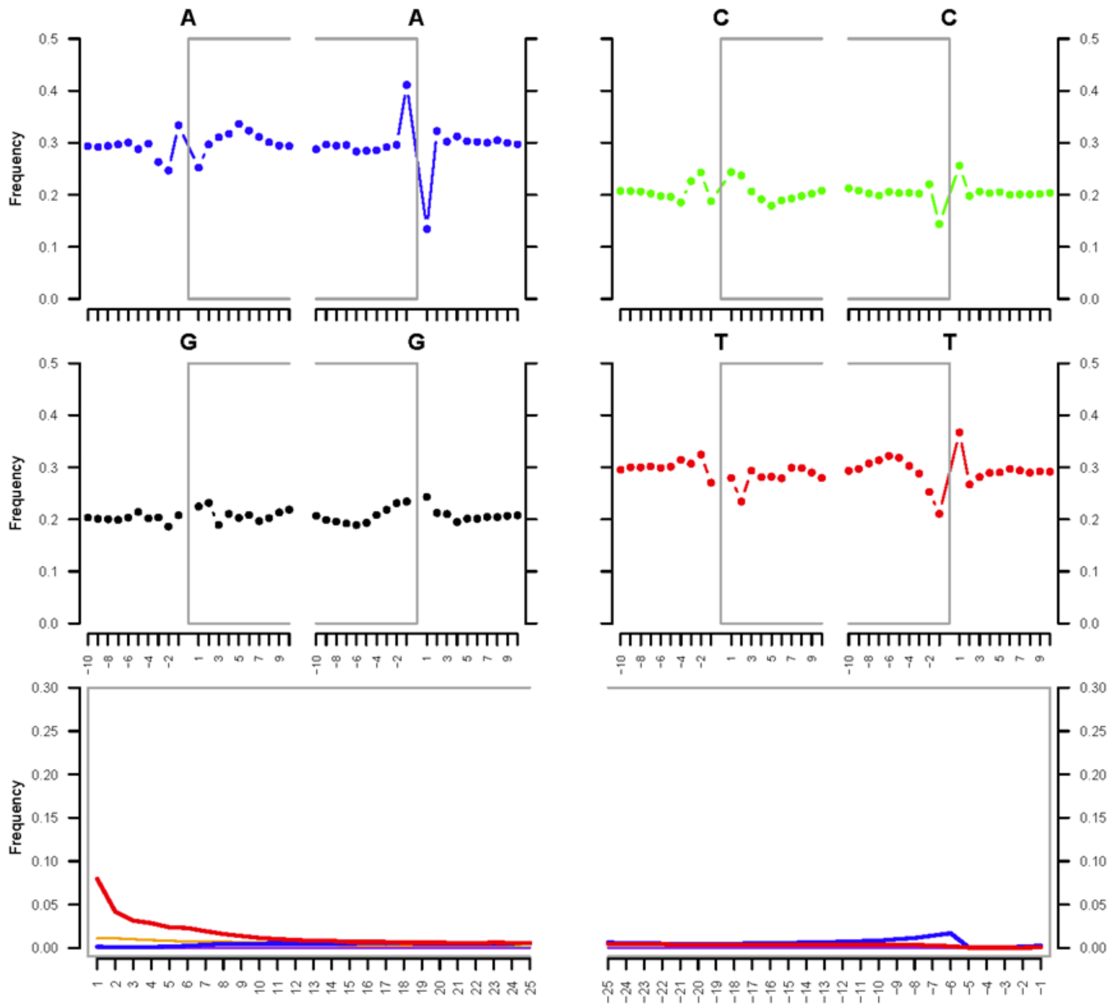


E. LD_34

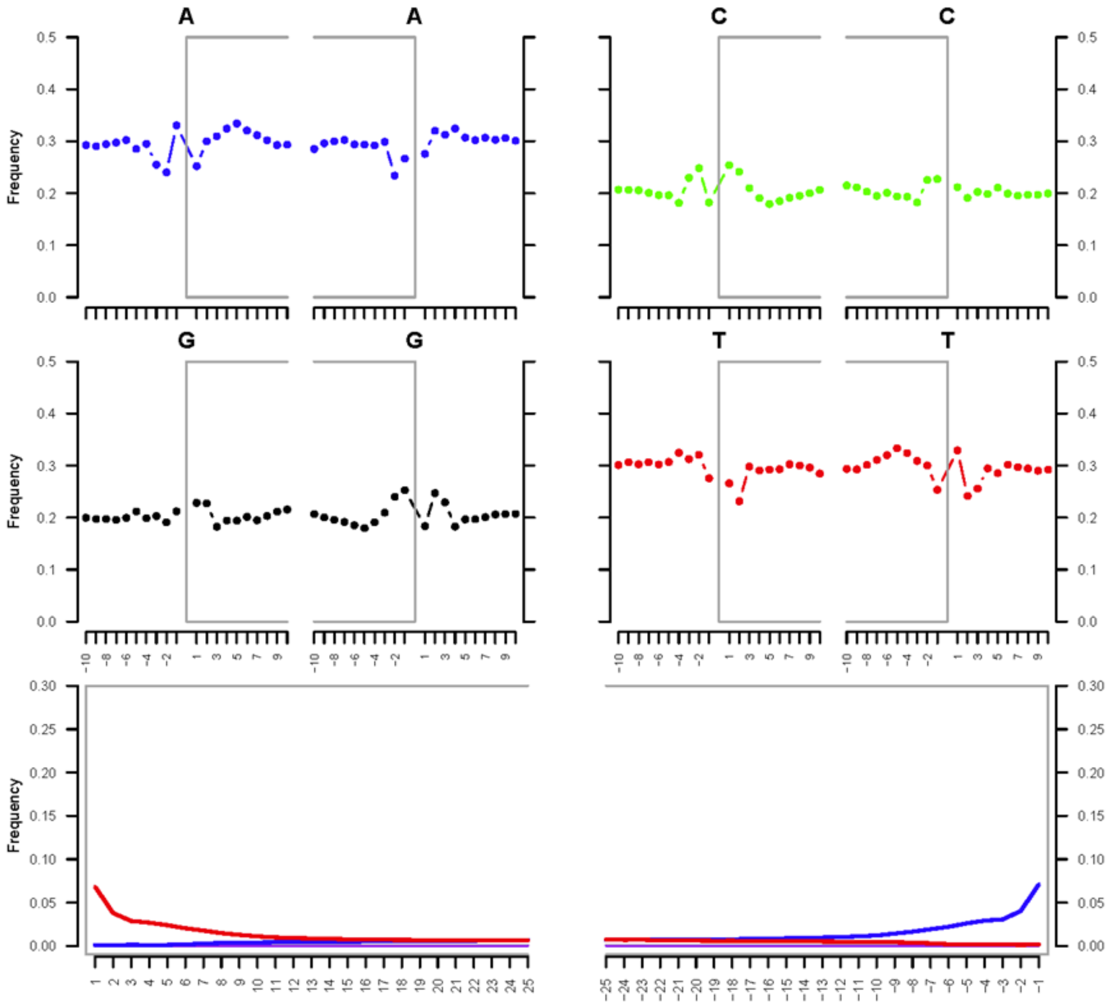


F. LD_36

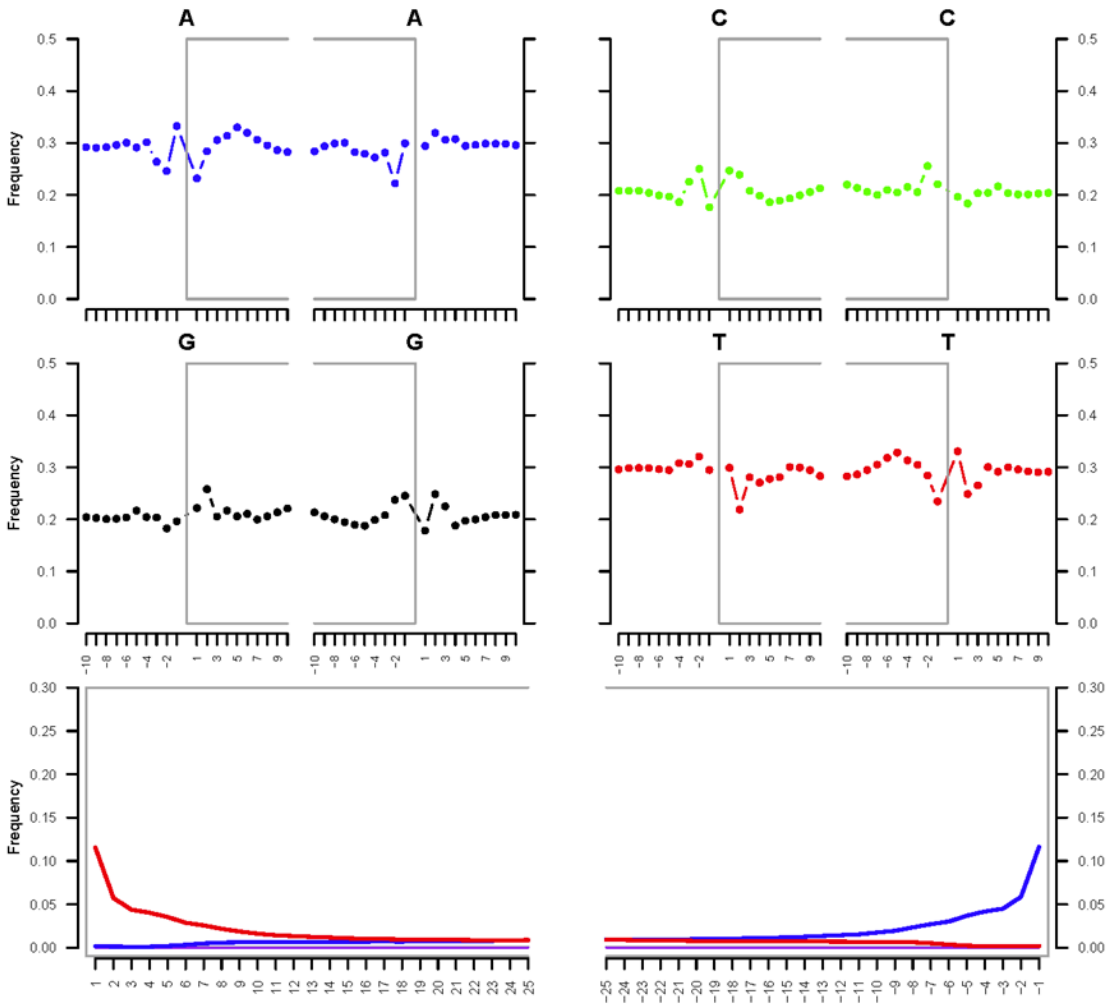


G. LD_38

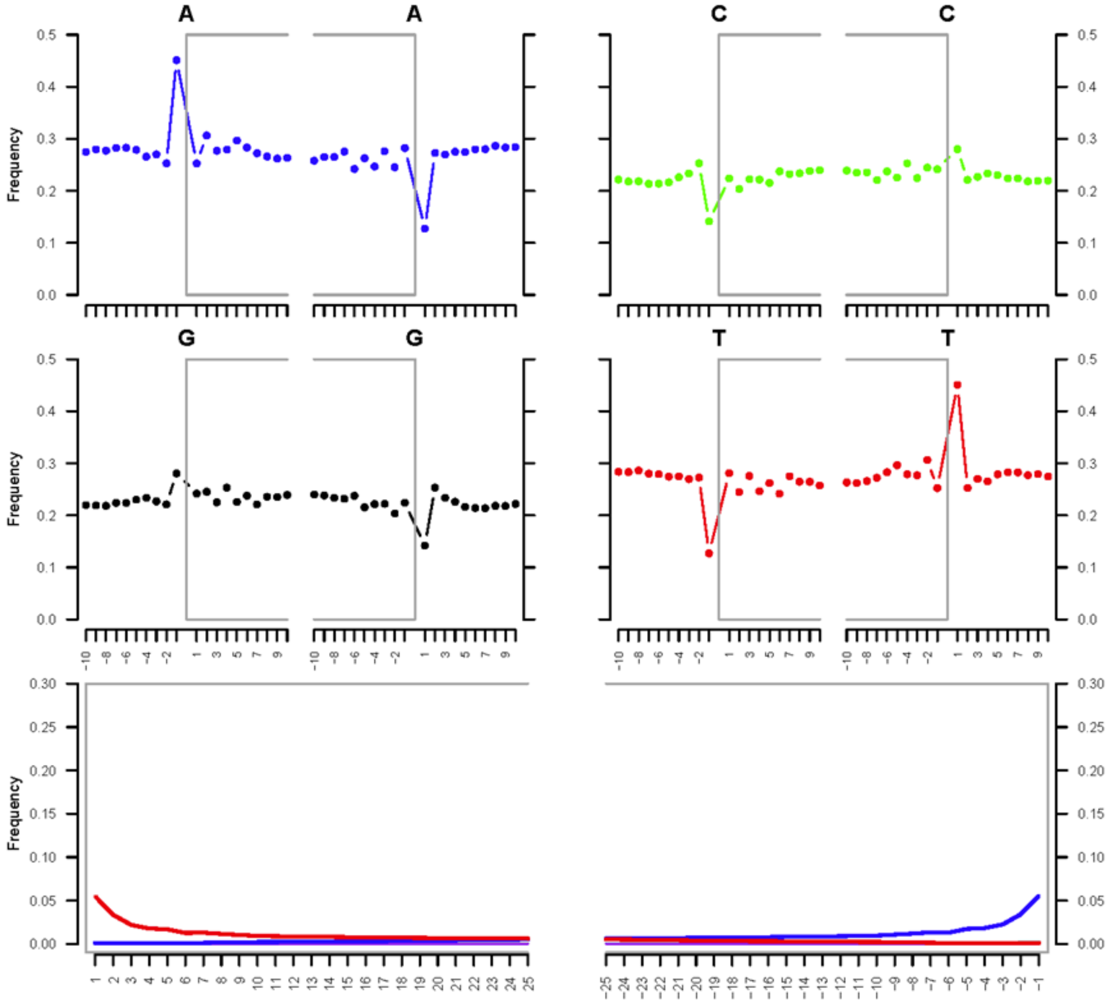


H. LD-TT

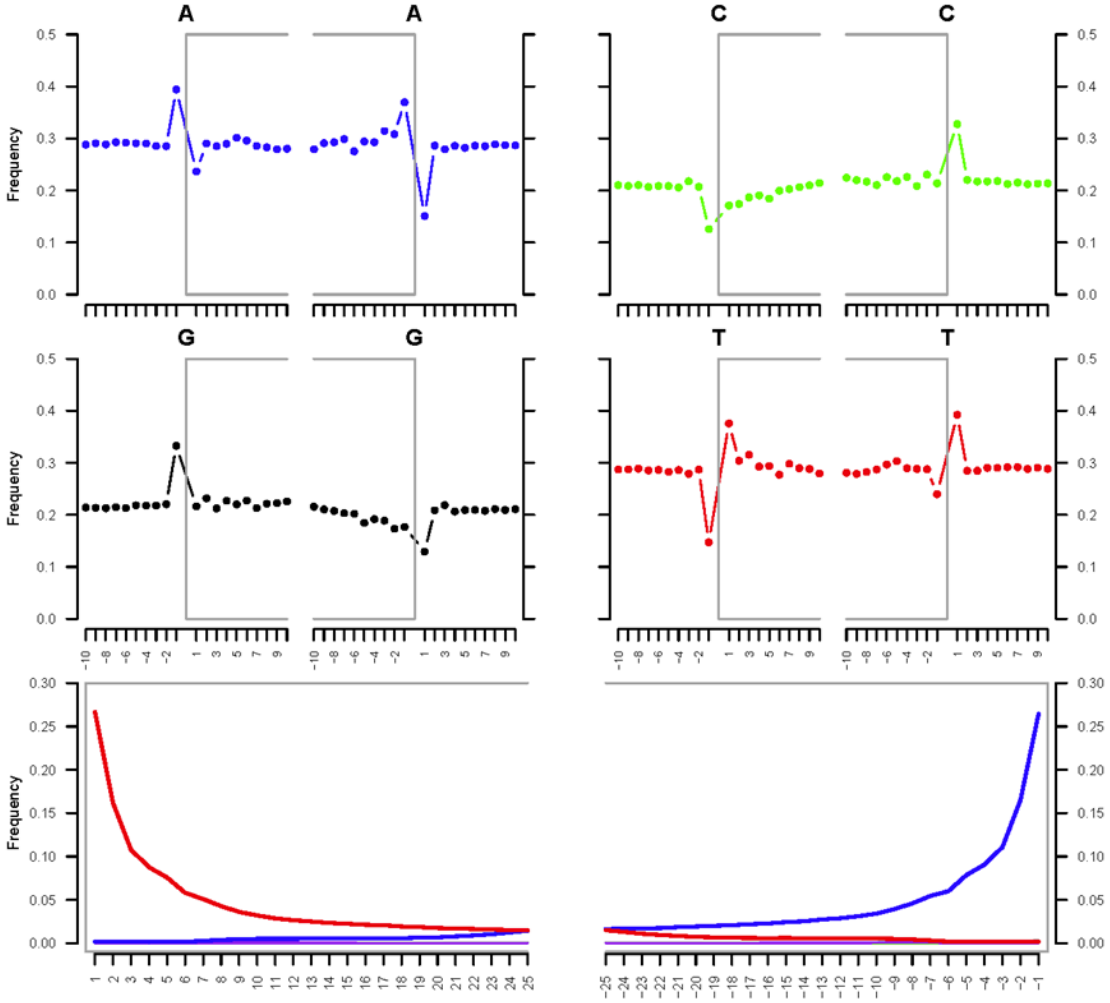


I. HANU-1

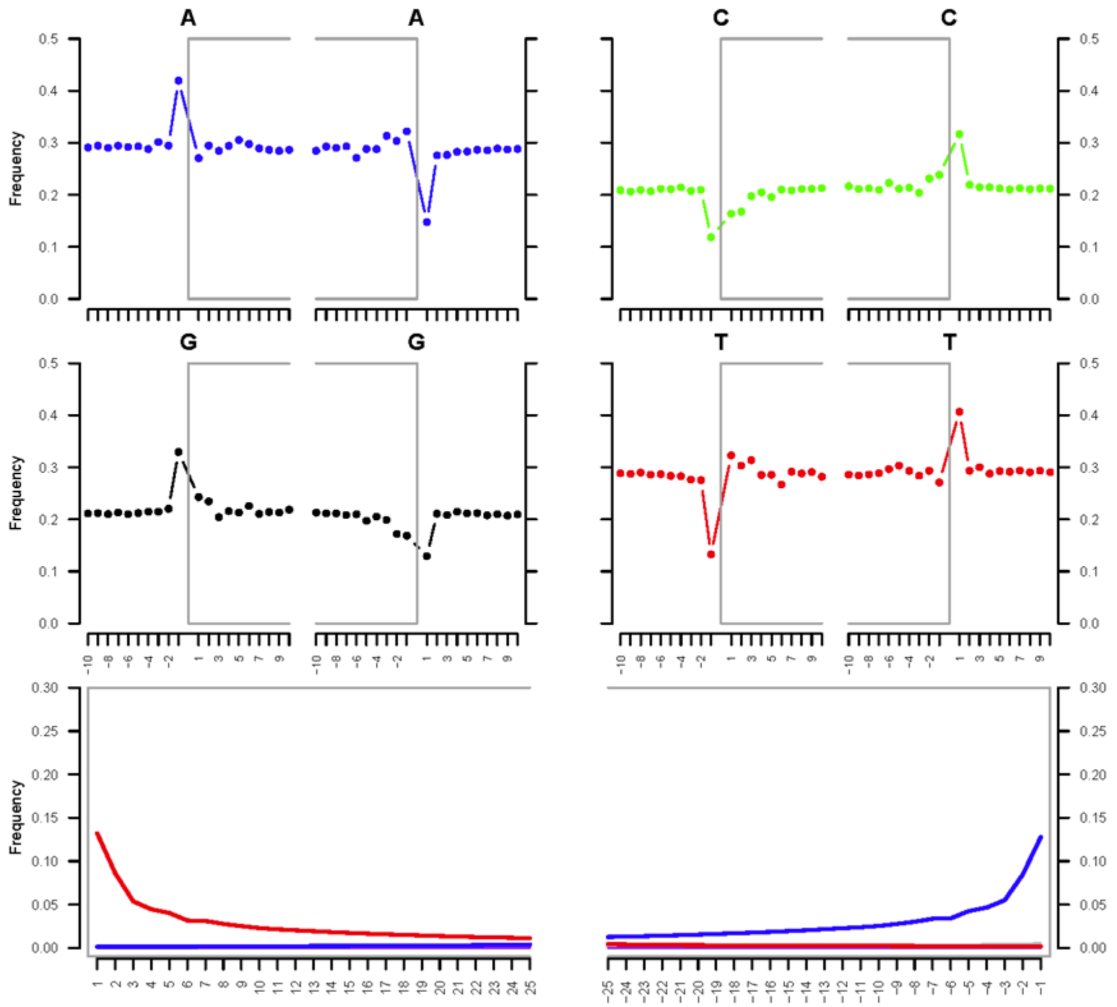


J. HANU-2

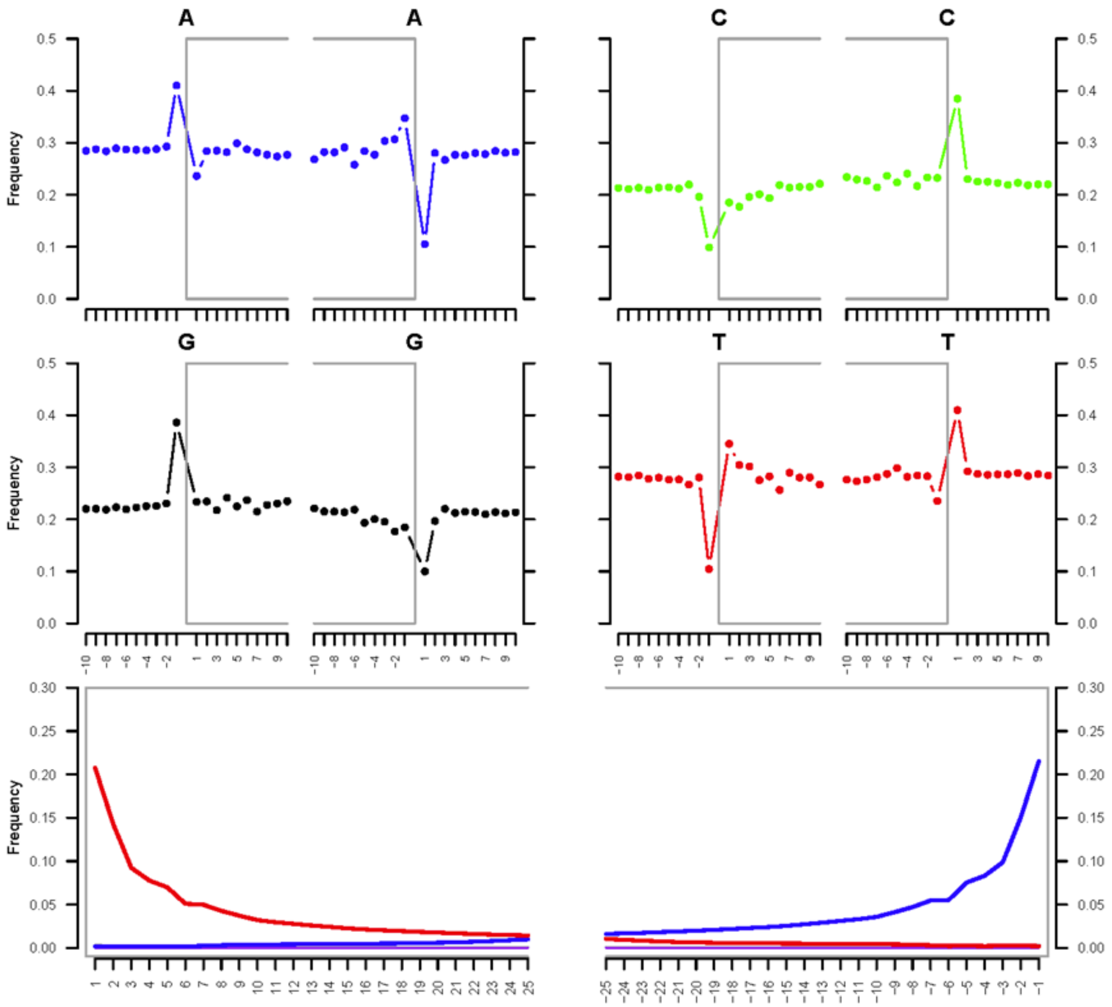


K. HANU-3

**Figure S1. (A-K) show mapDamage (fragmentation and misincorporation patterns) plots for the ancient Ladakh individuals generated using MapDamage v2.0.**

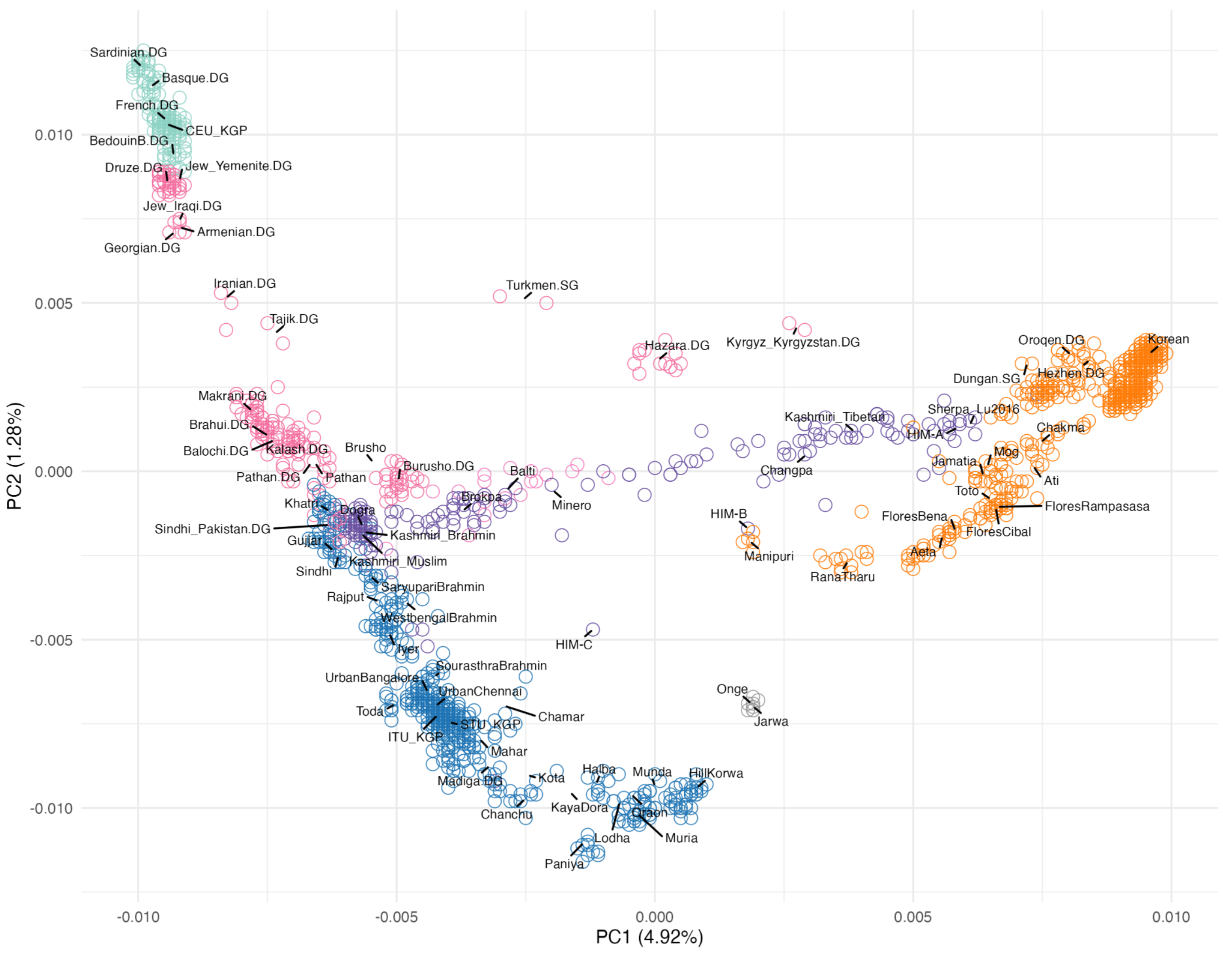


**Figure S2A. PCA of the reference space used in Figure 2A.** PCA showing only present-day populations used to compute PCs, spanning South Asia, the Tibetan Plateau, Central Asia, West Asia, East Asia, Southeast Asia, and Europe. This PCA is shown with population labels for the present-day references.

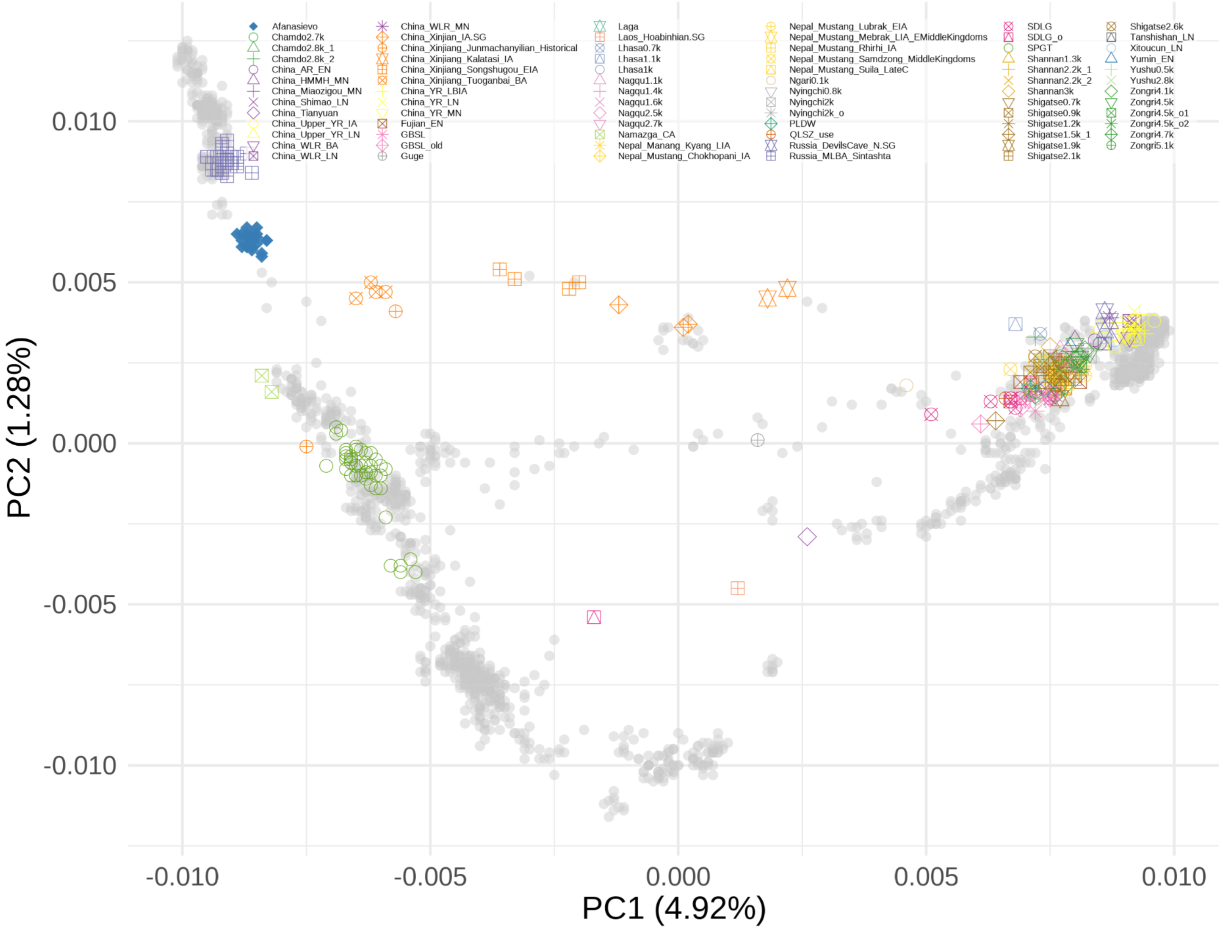


**Figure S2B. PCA of the reference space used in Figure 2A.** Select published ancient individuals from the Tibetan Plateau, East Asia, Central/South/West Asia, and Steppe were projected. Colored shapes represent ancient individuals based on archaeological, cultural or geographic affiliations as shown in the legend.

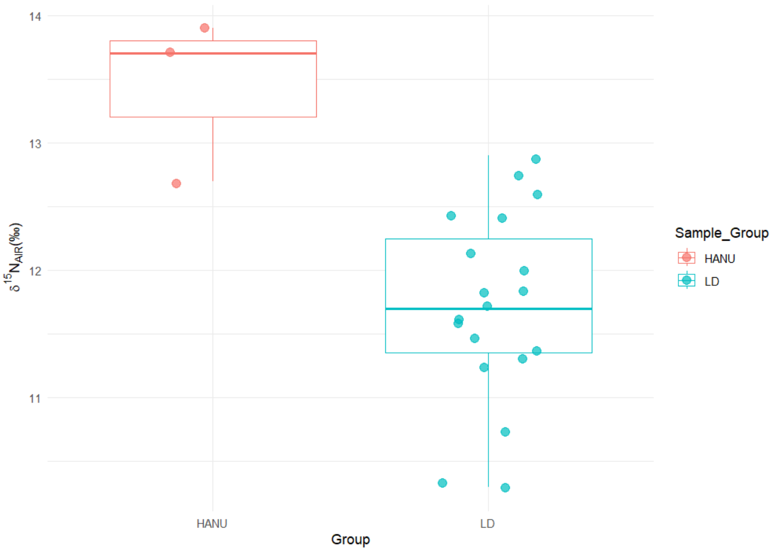


**A.**

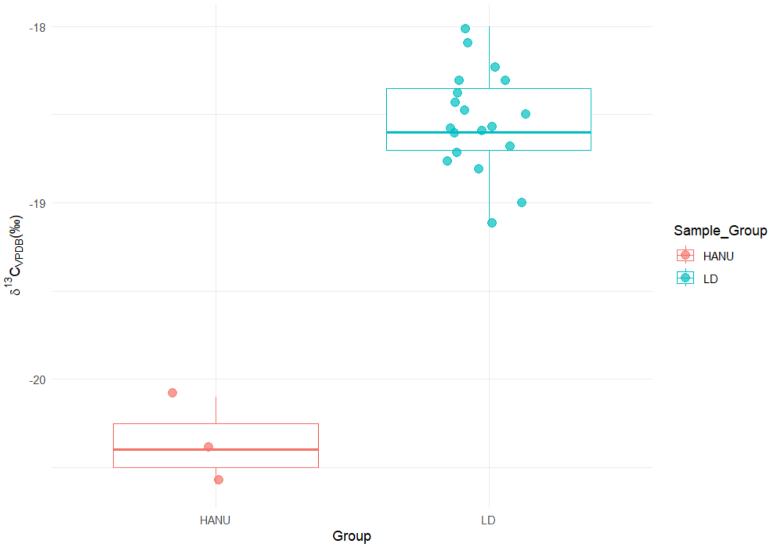


**B.**

**Fig. S3. Stable carbon and nitrogen isotopes for paleodietary reconstructions.** Boxplots representing A. $\delta^{15}N$ values (‰, AIR) and B. $\delta^{13}C$ values (‰, VPDB) for the ancient individuals from Hanu (HANU) and Old Lady Spider Cave (LD). $\delta^{15}N$ values are reported relative to the Ambient Inhalable Reservoir (AIR) standard and expressed in per mill (‰). $\delta^{13}C$ values are reported relative to the Vienna Pee Dee Belemnite (VPDB) standard and expressed in per mill (‰).

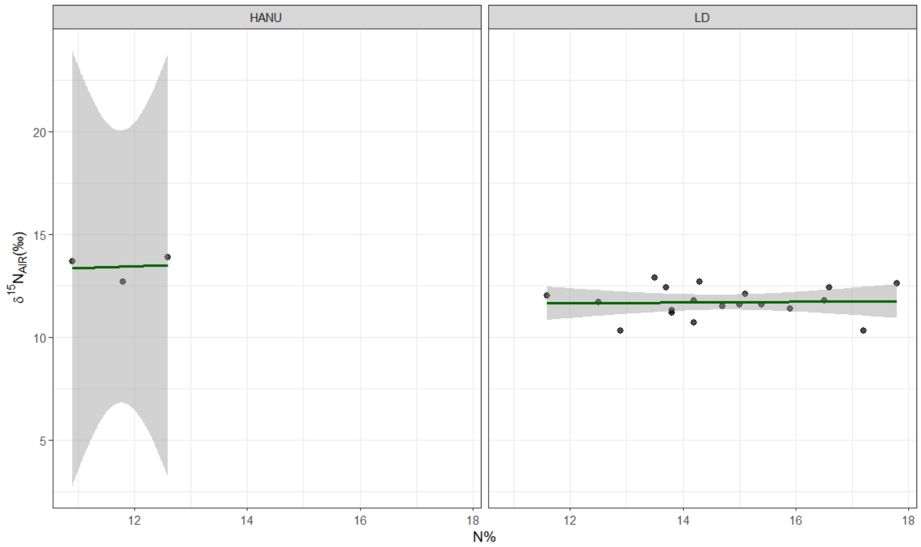


A.

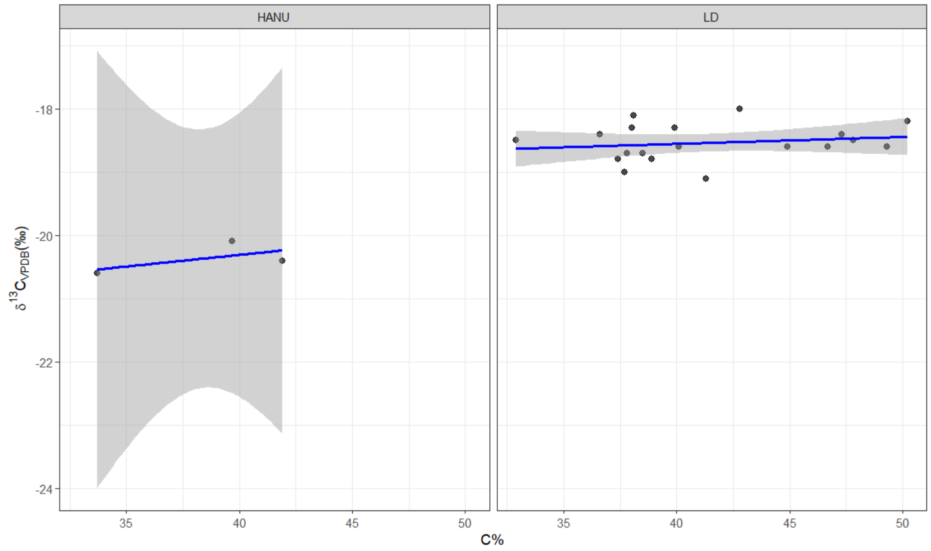


B.

**Fig. S4. Stable carbon and nitrogen isotopes for paleodietary reconstructions.** Scatter plots of stable isotope values versus elemental content in the ancient individuals from Hanu (HANU) and Old Lady Spider Cave (LD). A. $\delta^{15}N$ (‰, AIR) plotted against nitrogen content (%N), and B. $\delta^{13}C$ (‰, VPDB) plotted against carbon content (%C).

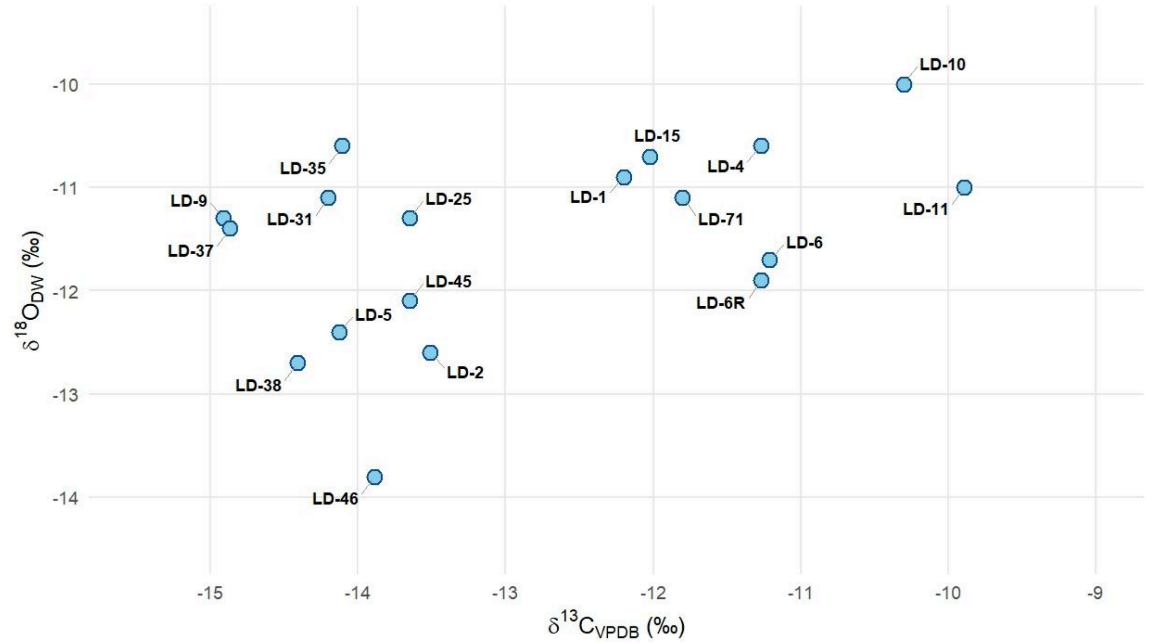


**Figure S5 Mobility patterns using oxygen and carbon isotopes.** Scatter plot of $\delta^{18}O_{dw}$ (‰) versus $\delta^{13}C$ (‰, VPDB) values measured in bone bioapatite from the ancient individuals from the Old Lady Spider Cave (LD). The $\delta^{18}O_{dw}$ (estimated drinking water) values are reported relative to the Vienna Standard Mean Ocean Water (VSMOW) standard and expressed in per mill (‰), here approximating the isotopic composition of ingested water. The $\delta^{13}C$ values are reported relative to the Vienna Pee Dee Belemnite (VPDB) standard and expressed in per mill (‰), reflecting the whole dietary carbon signal.

**Tables**

**Table S7A. Biological relatedness.** Pairwise relatedness estimates among Ladakh individuals inferred using BREADR.

| pair | relationship | pmr | sd | mismatch | nsnps | ave_rel | Same_Twins | First_ Degree | Second_ Degree | Unrelated |
|---|---|---|---|---|---|---|---|---|---|---|
| LD-01-LD-02 | Unrelated | 0.2456 | 0.0033 | 3998 | 16274 | 0.2502 | 0.00E+00 | 4.26E-73 | 9.03E-15 | 1.00E+00 |
| LD-01-LD-11 | Unrelated | 0.2498 | 0.0030 | 4927 | 19719 | 0.2502 | 0.00E+00 | 1.65E-101 | 6.85E-24 | 1.00E+00 |
| LD-01-LD-36 | First_Degree | 0.1893 | 0.0032 | 2763 | 14591 | 0.2502 | 3.39E-106 | 1 | 1.93E-17 | 1.15E-40 |
| LD-01-LD-TT | Unrelated | 0.2505 | 0.0034 | 3953 | 15776 | 0.2502 | 0.00E+00 | 3.93E-83 | 4.29E-20 | 1.00E+00 |
| LD-02-LD-11 | Unrelated | 0.2447 | 0.0033 | 4095 | 16731 | 0.2502 | 0 | 1.00E-72 | 4.80E-14 | 1.00E+00 |
| LD-02-LD-36 | Unrelated | 0.2513 | 0.0041 | 2759 | 10977 | 0.2502 | 1.76E-280 | 2.08E-59 | 7.81E-15 | 1.00E+00 |
| LD-02-LD-TT | Unrelated | 0.2556 | 0.0039 | 3114 | 12182 | 0.2502 | 0.00E+00 | 3.87E-74 | 2.72E-20 | 1.00E+00 |
| LD-11-LD-36 | Unrelated | 0.2493 | 0.0035 | 3735 | 14976 | 0.2502 | 0.00E+00 | 3.62E-76 | 8.44E-18 | 1.00E+00 |
| LD-11-LD-TT | Unrelated | 0.2528 | 0.0034 | 4095 | 16194 | 0.2502 | 0.00E+00 | 3.08E-91 | 2.12E-23 | 1.00E+00 |
| LD-36-LD-TT | Unrelated | 0.2528 | 0.0042 | 2657 | 10508 | 0.2502 | 2.34E-274 | 2.01E-59 | 2.03E-15 | 1 |

**Table S7B. Biological relatedness.** Pairwise relatedness inference using KIN.

| Pair | Relatedness | Second Guess | Log Likelihood Ratio | Within Degree Second Guess | Within Degree Log Likelihood Ratio | k0 | k1 | k2 | IBD Length | IBD Number |
|---|---|---|---|---|---|---|---|---|---|---|
| LD-01-LD-02 | Unrelated | Third Degree | 12.58 | | | 1 | 0 | 0 | 0 | 1 |
| LD-01-LD-11 | Unrelated | Third Degree | 17.434 | | | 1 | 0 | 0 | 0 | 1 |
| LD-01-LD-36 | Siblings | Second Degree | 13.583 | Parent-Child | 30.75148804 | 0.276 | 0.541 | 0.183 | 173 | 25 |
| LD-01-LD-TT | Unrelated | Third Degree | 9.242 | | | 0.945 | 0.055 | 0 | 11 | 4 |
| LD-02-LD-11 | Unrelated | Third Degree | 2.3 | | | 0.866 | 0.134 | 0 | 35 | 9 |
| LD-02-LD-36 | Unrelated | Third Degree | 7.942 | | | 0.948 | 0.052 | 0 | 10 | 2 |
| LD-02-LD-TT | Unrelated | Third Degree | 9.734 | | | 1 | 0 | 0 | 0 | 1 |
| LD-11-LD-36 | Unrelated | Third Degree | 13.985 | | | 1 | 0 | 0 | 0 | 1 |
| LD-11-LD-TT | Unrelated | Third Degree | 14.862 | | | 1 | 0 | 0 | 0 | 1 |
| LD-36-LD-TT | Unrelated | Third Degree | 9.843 | | | 1 | 0 | 0 | 0 | 1 |

**Table S8. Stable carbon and nitrogen isotopes for paleodietary reconstructions.** Bulk collagen stable isotope values of nitrogen ($\delta^{15}N$) and carbon ($\delta^{13}C$) and atomic C/N ratios for the individuals from the Old Lady Spider Cave (LD) and Hanu (HANU), Ladakh. $\delta^{15}N$ values are reported relative to the Ambient Inhalable Reservoir (AIR) standard, while $\delta^{13}C$ values are reported relative to the Vienna Pee Dee Belemnite (VPDB) standard. Both are expressed in parts per million (‰).

| Samples | $\delta^{15}N$ (‰, AIR) | Analysis | $\delta^{13}C$ (‰, VPDB) | Analysis | Atomic C/N |
|---|---|---|---|---|---|
| LD-01 | 11.8 | High animal protein | -18.4 | $C_3$ dominant | 2.9 |
| LD-02 | 12.4 | High animal protein | -18.5 | $C_3$ dominant | 2.9 |
| LD-05 | 12.1 | High animal protein | -18.3 | $C_3$ dominant | 2.9 |
| LD-07 | 11.5 | High animal protein | -18.8 | $C_3$ dominant | 3.1 |
| LD-08 | 12.4 | High animal protein | -18.4 | $C_3$ dominant | 3.1 |
| LD-09 | 10.7 | Moderate animal protein | -19 | Predominantly $C_3$ | 3.1 |
| LD-10 | 11.3 | High animal protein | -18.7 | $C_3$ dominant | 3.2 |
| LD-11 | 10.3 | Moderate animal protein | -18 | $C_3$ dominant | 3.3 |
| LD-14 | 11.8 | High animal protein | -18.3 | $C_3$ dominant | 3.3 |
| LD-17 | 11.2 | High animal protein | -18.8 | $C_3$ dominant | 3.2 |
| LD-18 | 12.7 | High animal protein | -18.1 | $C_3$ dominant | 3.1 |
| LD-25 | 11.7 | High animal protein | -18.6 | $C_3$ dominant | 3.2 |
| LD-34 | 10.3 | Moderate animal protein | -18.2 | $C_3$ dominant | 2.9 |
| LD-36 | 11.6 | High animal protein | -18.6 | $C_3$ dominant | 3.1 |
| LD-37 | 11.6 | High animal protein | -19.1 | $C_3$ dominant | 3.1 |
| LD-38 | 12.6 | High animal protein | -18.6 | $C_3$ dominant | 2.8 |
| LD-45 | 12 | High animal protein | -18.5 | $C_3$ dominant | 3.3 |
| LD-46 | 12.9 | High animal protein | -18.7 | $C_3$ dominant | 3.3 |
| LD-71 | 11.4 | High animal protein | -18.6 | $C_3$ dominant | 3.3 |
| LD-TT | 11.4 | High animal protein | -21.2 | $C_3$ diet | 3.1 |
| HANU-1 | 13.7 | High animal protein/Freshwater fish | -20.6 | $C_3$ diet | 3.1 |
| HANU-2 | 12.7 | High animal protein | -20.1 | $C_3$ diet | 3.4 |
| HANU-3 | 13.9 | High animal protein/Freshwater fish | -20.4 | $C_3$ diet | 3.3 |

**Table S9. Stable carbon and oxygen isotopes for late-life mobility reconstructions.** Carbon ($\delta^{13}C$) and Oxygen ($\delta^{18}O$) stable isotopic values with standard deviations (SD), and derived estimated drinking water oxygen isotopic values ($\delta^{18}O_{dw}$) from archaeological site of the Old Lady Spider Cave (LD) of Ladakh. The $\delta^{13}C$ values are reported relative to the Vienna Pee Dee Belemnite (VPDB) standard and expressed in parts per million (‰). The $\delta^{18}O$ values are reported relative to both Vienna Pee Dee Belemnite (VPDB) standard and Vienna Standard Mean Ocean Water (VSMOW) standard and expressed in parts per million (‰). All individuals are classified local or non-local on the basis of $\delta^{18}O_{dw}$ values. See methods for details on conversion between VPDB and VSMOW and the estimation of $\delta^{18}O_{dw}$.

| Samples | $\delta^{13}C$ (‰, VPDB) | $\delta^{13}C$ SD | Analysis | $\delta^{18}O$ (‰, VPDB) | $\delta^{18}O$ SD | $\delta^{18}O$ (‰, VSMOW) | Estimated $\delta^{18}O$ (‰, dw) | Local/ Non-Local |
|---|---|---|---|---|---|---|---|---|
| LD-01 | -12.2 | 0.04 | $C_3$ | -10.92 | 0.03 | 19.7 | -10.9 | Local |
| LD-02 | -13.51 | 0.07 | $C_3$ | -12.18 | 0.04 | 18.4 | -12.6 | Local |
| LD-04 | -11.27 | 0.07 | $C_3$ | -10.66 | 0.07 | 19.9 | -10.6 | Local |
| LD-05 | -14.13 | 0.04 | $C_3$ | -11.97 | 0.06 | 18.6 | -12.4 | Local |
| LD-06 | -11.21 | 0.04 | $C_3$ | -11.48 | 0.03 | 19.1 | -11.7 | Local |
| LD-06R | -11.27 | 0.06 | $C_3$ | -11.62 | 0.03 | 18.9 | -11.9 | Local |
| LD-09 | -14.91 | 0.05 | $C_3$ | -11.19 | 0.12 | 19.4 | -11.3 | Local |
| LD-10 | -10.3 | 0.07 | $C_3$ dominant | -10.23 | 0.03 | 20.4 | -10 | Local |
| LD-11 | -9.89 | 0.05 | $C_3$ dominant | -10.97 | 0.11 | 19.6 | -11 | Local |
| LD-15 | -12.02 | 0.04 | $C_3$ | -10.73 | 0.03 | 19.9 | -10.7 | Local |
| LD-25 | -13.65 | 0 | $C_3$ | -11.15 | 0.03 | 19.4 | -11.3 | Local |
| LD-31 | -14.2 | 0.03 | $C_3$ | -11.04 | 0.09 | 19.5 | -11.1 | Local |
| LD-35 | -14.11 | 0.03 | $C_3$ | -10.64 | 0.1 | 19.9 | -10.6 | Local |
| LD-37 | -14.87 | 0.1 | $C_3$ | -11.27 | 0.13 | 19.3 | -11.4 | Local |
| LD-38 | -14.41 | 0.07 | $C_3$ | -12.22 | 0.16 | 18.3 | -12.7 | Local |
| LD-45 | -13.65 | 0.06 | $C_3$ | -11.78 | 0.07 | 18.8 | -12.1 | Local |
| LD-46 | -13.89 | 0.07 | $C_3$ | -13.04 | 0.02 | 17.5 | -13.8 | Local |
| LD-71 | -11.8 | 0.04 | $C_3$ | -11.04 | 0.04 | 19.5 | -11.1 | Local |
